# Identical Sequences, Different Behaviors: Protein Diversity Captured at the Single-Molecule Level

**DOI:** 10.1101/2021.02.24.432730

**Authors:** Rafael Tapia-Rojo, Alvaro Alonso-Caballero, Carmen L. Badilla, Julio M. Fernandez

## Abstract

The classical “one sequence, one structure, one function” paradigm has shaped much of our intuition of how proteins work inside the cell. Partially due to the insight provided by bulk biochemical assays, individual biomolecules are assumed to behave as identical entities, and their characterization relies on ensemble averages that flatten any conformational diversity into a unique phenotype. While the emergence of single-molecule techniques opened the gates to interrogating individual molecules, technical shortcomings typically limit the duration of these measurements to a few minutes, which prevents to completely characterize a protein individual and, hence, to capture the heterogeneity among molecular populations. Here, we introduce a magnetic tweezers design, which showcases enhanced stability and resolution that allows us to measure the folding dynamics of a single protein during several uninterrupted days with a high temporal and spatial resolution. Thanks to this instrumental development, we do a complete characterization of two proteins with a very different force-response: the talin R3^IVVI^ domain and protein L. Days-long recordings on the same single molecule accumulate several thousands of folding transitions sampled with sub-ms resolution, which allows us to reconstruct their free energy landscapes and describe how they evolve with force. By mapping the nanomechanical identity of many different protein individuals, we directly capture their molecular diversity as a quantifiable dispersion on their force response and folding kinetics. Our instrumental development offers a new tool for profiling individual molecules, opening the gates to the characterization of biomolecular heterogeneity.

## Introduction

Traditionally, it is assumed that a protein’s sequence univocally dictates its structure, which consequently determines its function. ^1^ For example, the classical funnel model for protein folding offers an intuitive vision of how the myriad of energetically-disfavored conformations accessible to a given residue sequence eventually collapse to a unique free energy basin that defines the functional native state. ^2,3^ Bulk biochemical and structural techniques have supported this paradigm, as the experimental signal is an intrinsic ensemble average over a population of thermodynamic size. However, increasing evidence resulting from novel experimental approaches have challenged this dogma, suggesting that proteins are dynamic entities with an intrinsic diversity arising from more complex conformational landscapes.^4–8^

In this context, the emergence of single-molecule force spectroscopy techniques shifted the paradigm of biophysical experimentation by allowing to interrogate individual molecules under force to directly sample the distributions of their molecular properties without relying on pre-averaged signals.^9,10^ While protein nanomechanics has been typically approached using atomic force miscroscopy (AFM) and optical tweezers, these two techniques suffer from large mechanical drift that limits the measurements to a few minutes, eventually requiring to interrogate several molecular samples to accumulate enough statistics.^11,12^ By contrast, magnetic tweezers, an intrinsically more stable technique, has been mostly devoted to the study of nucleic acids, likely owing to the lower temporal and spatial resolution of the early instrumental designs, and the initial difficulty to obtain specific and long lasting tethers.^10,12–16^ However, recent instrumental advances and the development of novel anchoring strategies are now demonstrating the expediency of magnetic tweezers for measuring protein mechanics.^17–20^ Magnetic tweezers have the crucial advantage of offering intrinsic force-clamp conditions, which afford direct control of the intensive variable (force), while the extensive variable (extension) is measured, providing the natural statistical ensemble for studying protein dynamics in equilibrium. In this sense, by pushing the instrumental boundaries to increase the stability and resolution limitations, magnetic tweezers could enable to fully portray the nanomechanical properties of a single protein and, hence, open the gates to capturing the diversity of protein phenotypes and directly characterizing their functional heterogeneity.

Here, we present a novel magnetic tweezers design that enables us to do a full nanomechanical characterization of a single protein through long-term ultra-stable recordings at high temporal and spatial resolution. Our setup highlights its simple and bespoke design from off-the-shelf and affordable components that make it a highly accessible instrument, which we describe with all details necessary for its installation in any lab in the world. Thanks to this instrumental development, we measure days-long recordings of the folding dynamics of two different proteins, the talin R3^IVVI^ domain and protein L, obtaining a complete nanomechanical mapping using a single molecular individual on each case. Our data demonstrates that polymer physics is a dominant component of protein folding under force, which governs the refolding transition by dictating the shape of the unfolded basin and the evolution of the free energy landscape with force. To directly explore the nanomechanical diversity of these two proteins, we measure several protein individuals and characterize the force response of each molecule, which shows an extent of dispersion both on its folding equilibrium and kinetics while their elastic properties remain homogeneous, validating our calibration and fingerprinting. Our results strongly suggest that proteins have heterogeneous nanomechanical properties, previously veiled by the limited stability of the available techniques, and that could result in a spectrum of functional heterogeneity.

## METHODS

### Instrumental Design

In magnetic tweezers force spectroscopy, a single molecule is immobilized between a glass cover slide and a superparamagnetic bead. By controlling a magnetic field gradient, the molecule is subjected to pN-level forces, while its extension is measured by video-tracking the bead diffraction or interference patterns, depending on the illumination geometry.^17,18,21^ Due to the overly compliant trap created by the magnetic field, magnetic tweezers offers passive force-clamp conditions that afford direct control of the pulling force, which permits to interrogate single molecules in equilibrium.

We present here a novel magnetic tweezers setup, assembled from off-the-shelf and affordable components. Figure 1 shows a three dimensional model of our setup, indicating each of its elements (see SI Appendix for the parts list). Our setup has a simple and dedicated microscope design, using an optical arrangement for a 160-mm objective (see Fig. S1 and SI Appendix for the description and schematics of the optical pathway). Briefly, we illuminate from below using a LED source (Thorlabs, element #12), and collimate the light with a plano-convex lens (Thorlabs, element #10) to focus it on the objective with a tube lens (Thorlabs, element #7), while the image is recorded with a CMOS camera (Ximea, element #14).

**Figure 1:**
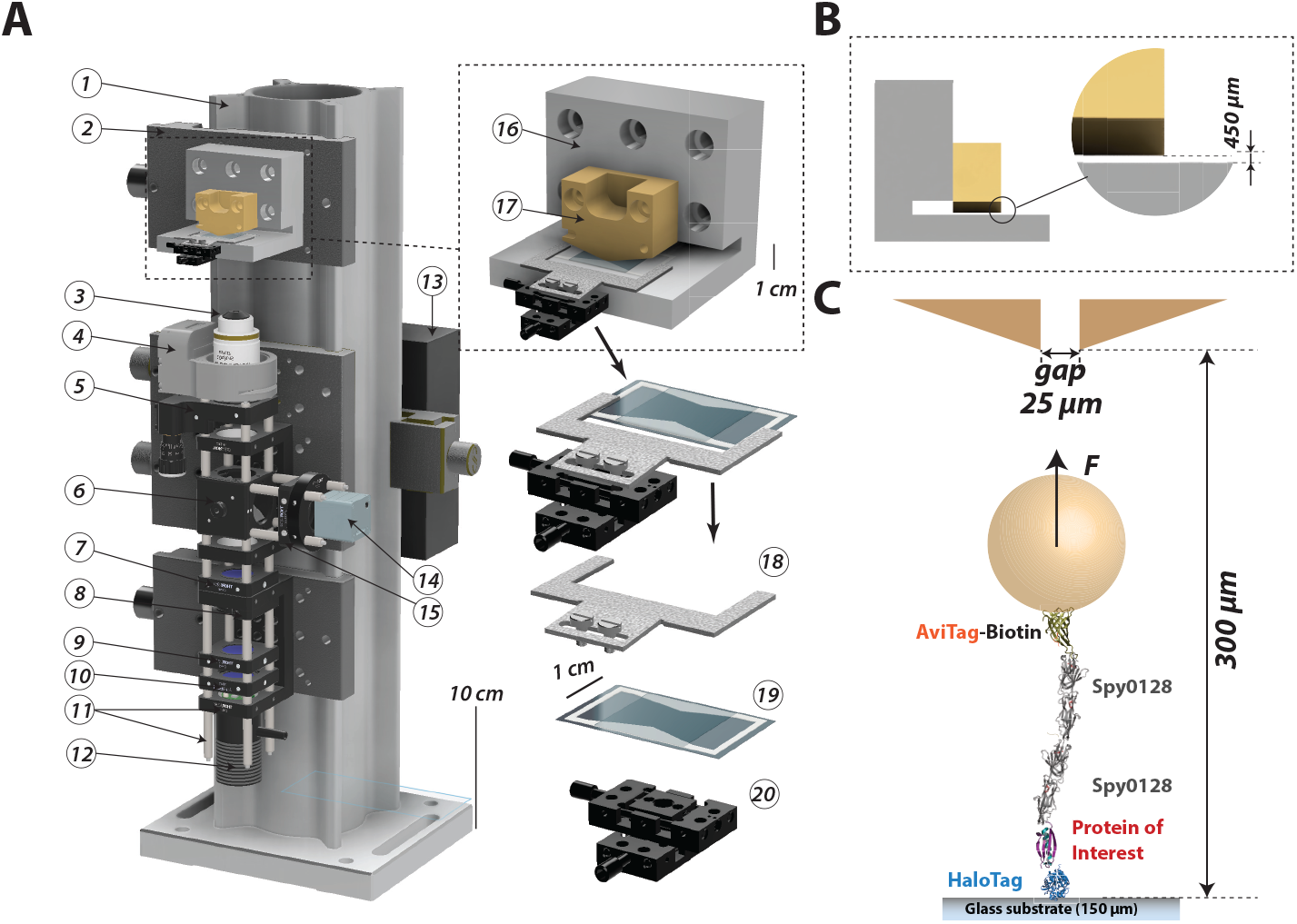
Three-dimensional model of the magnetic tweezers setup: (A) The setup comprises the following components: (1) 95-mm construction rail; (2) optical rail carriage; (3) 100x 160 mm objective; (4) piezo scanner; (5) Z flexor stage; (6) 50:50 beam splitter; (7) tube lens; (8) diaphragm; (9) filter; (10) plano-convex lens; (11) cage-rod system; (12) LED light source; (13) electronics box; (14) CMOS camera; (15) plano-convex lens; (16) mount piece (card-reader); (17) magnetic tape head; (18) manipulation fork; (19) flow cell; (20) XY linear stage. (B) Detail of the card reader piece. The tape head is maintained at a distance of 450 *μ*m, which, when using 150 *μ*m thick bottom glasses, positions the gap 300 *μ*m away from the magnetic beads. (C) Schematics of our anchoring strategy for single-protein magnetic tweezers experiments. The protein of interest is inserted between a HaloTag and two Spy0128 repeats, following a biotilinated AviTag for tethering with streptavidin-coated superparamagnetic beads (M-270). This strategy provides a highly specific and stable anchor and an inextensible molecular handle.

While most magnetic tweezers designs control the force by displacing a pair of magnets using a motor or a voice coil,^17,18^ we use here a magnetic tape head (Brush Industries, element #17), an old piece of technology more commonly employed to read magnetic bands in music cassettes or credit cards.^19^ This implementation offers enhanced control of the pulling force through the electric current supplied to the tape head, which can be modulated with high accuracy and speed that enables us to do swift force changes or to apply complex force signals such as mechanical noise or force oscillations.^19,22^ To generate calibrated and reproducible forces, it is paramount to position the tape head over the magnetic probe with micron precision, given that the magnetic field generated by the tape head changes over a length scale defined by the gap width, 25 *μ*m in our case. To solve this problem, we introduce here a mount piece fabricated with high-precision CNC machining (element #16, see Fig. S2)—dubbed the “card-reader”—where we mount the tape head with a dowel-pin system. Thanks to the card-reader, the tape head gap is positioned precisely at 450 *μ*m over the surface so that, when using 150 *μ*m thick bottom glasses, the gap is at 300 *μ*m over the magnetic probe (Fig. 1B). The tape head is maintained under electronic feedback with a PID current-clamp circuit, which showcases a bandwidth greater than 10 kHz (see Fig. S3 and SI Appendix). Importantly, the electronic circuit and the tape head is powered by a pair of 12 V batteries—lead-acid 18 Ah motorbike batteries—which provide a clean and stable source of electric current, key to conduct swift force changes at very high rates.

The introduction of the card-reader piece is of capital importance for the ultra-stable conditions of our magnetic tweezers design, allowing us to apply highly reproducible and precise forces by simply controlling the electric current supplied to the tape head as:

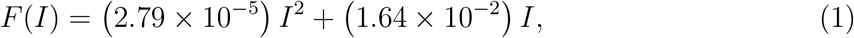

where *F* is the pulling force in pN and *I* the electric current in mA. As described in, ^19^ the parameters in Eq. (1) are determined by the specifics of the tape head arrangement, including the vertical distance separating the gap and the bead, and the magnetic properties of the superparamagnetic beads, the M-270 Dynabeads in the present case. In our implementation, we can apply forces between 0 and 44 pN when using electric currents between 0 and 1,000 mA, and do force changes as small as ~10^−2^ pN through electric current changes of ~1 mA. The force range can be increased by using magnetic beads of a larger diameter or higher magnetization. For instance, the implementation of the M-450 Dynabeads increases the maximum force to ~220 pN.^23^

The simplified dedicated microscope of this magnetic tweezers setup, combined with the capability of applying precise constant forces with the card-reader piece, results in an ultrastable instrument compared to designs that implement standard microscopes, which permits doing continuous measurements during several days with negligible drift. In the SI Appendix, we include a practical guide with all details for reproducing our setup, including the parts list, electronic circuit diagram, and technical drawing for the card-reader piece.

### Fluid Chamber Preparation

We conduct our single-molecule experiments in single-use fluid chambers (element #19) built by sandwiching two glass cover-slides (Ted Pella) with a custom-designed laser-cut parafilm pattern. Using a 40×24 mm bottom glass and a 22×22 mm top glass, the fluid chambers shows two open wells that allow for buffer exchanges, useful, for instance, for altering in real-time the chemical conditions of the experiment.^24,25^ Additionally, when conducting long-term experiments, the fluid chamber can be sealed using high-density white oil, which greatly stabilizes the buffer conditions and prevents evaporation. The bottom glass is functionalized to harbor the HaloTag O4 ligand, as previously described,^17^ while the top glass is treated with repel silane to hydrophobize it and allow solution flow inside the chamber (see Fig. S4 and SI Appendix). The fluid chamber is clipped on a metal-stamped manipulation fork (element #18) using double-sided tape. The fork is mounted on an XY-stage (Newport, element #20) using two high-precision shoulder screws. The XY-stage is used to scout for beads over small 3×3 mm areas, but the fork can be loosened to displace it laterally and move to new regions in the fluid cell.

### Molecular construct for single-molecule studies

To measure individual protein domains during extended periods of time, it is necessary to develop a long-lasting anchoring technique and molecular handles that interfere minimally with the signal of the protein of interest. Our strategy implements HaloTag chemistry for covalent and specific anchoring of the protein construct to functionalized glass cover slides, as previously discussed,,^11,17^ while the other terminus is tethered to streptavidin-coated beads through a biotinilated AviTag (Fig. 1C). To measure single protein domains, we have here designed a molecular template comprising two repeats of the Spy0128 protein, which is inextensible under force owing to the isopeptide bonds that clamp its termini (Fig. 1C).^26,27^ In this simple anchoring strategy, the protein of interest is inserted between the HaloTag enzyme at the N-terminus and the Spy0128 repeats, which provides a ~15 nm-long stiff molecular handle that shows no mechanical response, hence, allowing the direct measurement of the dynamics of the protein of interest. While the biotin-streptavidin tethering at the C-terminus is stable enough to withstand typical folding forces (<15 pN) during extended periods of time, we have developed more enduring strategies that involve HaloTag anchors at both termini, which permit to pull at arbitrarily high forces.^23,25^ We include in the SI Appendix (Fig. S5) a detailed vector map for cloning and detailed protocols for protein cloning, expression, and purification.

### Data Acquisition and Analysis

We have developed a custom-written software in C++/Qt for data acquisition. Briefly, at the start of an experiment, we build a z-stack library of the magnetic and reference beads taking 20 nm steps using the piezoelectric actuator with a total of 128 images. Then, the radial vector of each image is calculated from the Fourier Transform with a pixel-addressing algorithm.^17^ In the course of an experiment, the z-extension of the molecule is computed in real time by correlating the radial vectors of the magnetic and reference beads with the stack library and calculating the optimal Gaussian fit of the Pearson correlation coefficients using Caruana’s algorithm.^28^ The use of an analytical method instead of a minimum-square fit greatly accelerates the image processing algorithm, which, with our current implementation, can reach sampling rates of ~1.7 kHz (~580 *μ*s). The software writes in real time a sequential file containing the time mark, protein extension, and force stored in binary format. The file can be accessed using an independent software (we employ mostly Igor Pro 8, Wavemetrics) to visualize in real-time the course of the measurement without interfering with the data acquisition process. We include as part of our SI Appendix the source code for the data acquisition, which has been uploaded to the online repository GitHub.

The combination of stability and the high sampling density of our instrument results in large volumes of data, which, for instance, can easily reach ~6×10^8^ data points for a single protein measured over a few days. Such large volume of data can be arduous to handle and analyze manually, raising the same kind of problems that the Big Data field deals with. To this aim, we have developed a JSON-based file format for single-molecule data, which enables the direct access to specific experiment fragments for its automatic analysis. JSON (JavaScript Open Notation) is an open source file format compatible with most programming languages and software—including Python, C++, Matlab, or Igor Pro— that organizes the data by attribute-value pairs or “keys”, which enables to automatically load these data portions instead of sequentially accessing the whole file. In our case, the architecture of our file format uses keys defined by the pulse protocols employed to measure the molecule following a “F/T/flag” format, where F is the employed force in pN, T the duration of the pulse in seconds, and flag either “C” for constant force, or “L” for a linear force ramp. Under a protocol key, the data is stored hierarchically as arrays identified by their key, “time” (time mark in s), “z” (protein extension in nm), and “force” (applied force in pN). Figure 2A shows a representation of our JSON-based file format, which illustrates its architecture. With this implementation, days-long experiments are stored in a rational format, which allows processing large amounts of data in an automatic way and with minimal human intervention.

**Figure 2:**
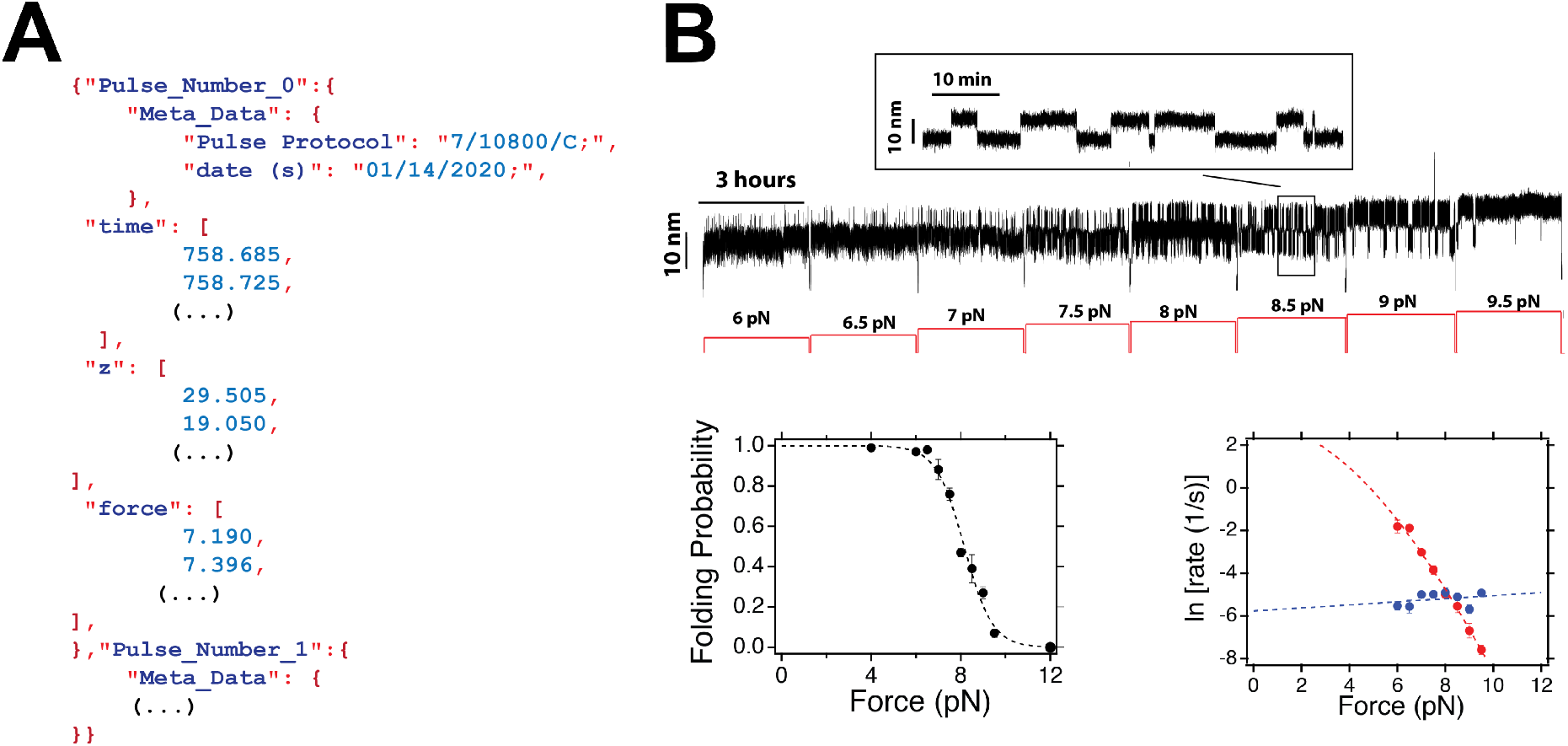
A JSON-based file format for automatic single-protein data analysis: (A) Architecture of the JSON-based single-molecule data format. The file is structured in attribute-values (“keys”) according to the experimental force-pulse protocol. Each attribute contains the data as arrays with the time mark (“time”), protein extension (“z”), and force (“force”). (B) Force dependency of protein L folding dynamics measured through JSON-pulse protocols. One day-long recording, exploring different forces for 3 hours, following JSON-based pulses (upper panel). Given the rational organization of the data based on the utilized experimental protocol, it is possible to automatically access and analyze each fragment following the JSON keys. The folding probability and folding (red) and unfolding (blue) rates are automatically analyzed from the JSON file format (lower panel). This strategy enables the manipulation and analysis of large data sets with minimal human intervention.

As an illustration of the potential of the JSON format approach, Fig. 2B shows the fragment of a protein L recording, aimed at directly characterizing the force-dependency of its folding dynamics. This 24 hours-long recording is organized in eight 3 hours-long JSON pulses, each exploring a different force between 6 and 9.5 pN, covering the full range over which protein L undergoes reversible folding dynamics. By storing this recording in our JSON-file format, the experimental data can be automatically accessed and analyzed according to the employed pulse, in this case the explored force. Hence, the folding properties of protein L, portrayed by its folding equilibrium and folding kinetics (Fig. 2B, lower panel), can be automatically calculated by simply analyzing each pulse fragment.

## Results

Our experimental technique permits the continuous measurement of protein folding dynamics during extended timescales and at high resolution. To illustrate our approach, we characterize the folding dynamics of the talin R3^IVVI^ and protein L using a single molecular individual. This allows us to compare the nanomechanical performance of different molecular specimens and directly observe the extent of diversity between their nanomechanical performances.

### Nanomechanics of the Talin R3^IVVI^ domain

To characterize the nanomechanics of the R3^IVVI^ domain, we collect a single 27 hour-long recording on the same single protein, employing the molecular construct discussed above (Fig. 3A). The fast folding dynamics of this protein allow accumulating over 70,000 folding and unfolding transitions along the whole experiment, and fully characterize the force-dependence of its folding equilibrium. Figure 3B shows 50 seconds-long fragments of R3^IVVI^ folding dynamics at three different forces. These transitions between the folded (low extension) and unfolded (high extension) states yield ~20 nm changes on the protein extension, which gives rise to the characteristic “hopping” dynamics. As displayed in Fig. 3B, R3^IVVI^ is highly sensitive to the pulling force, and changes of a fraction of a pN are sufficient to shift its folding equilibrium considerably.

**Figure 3:**
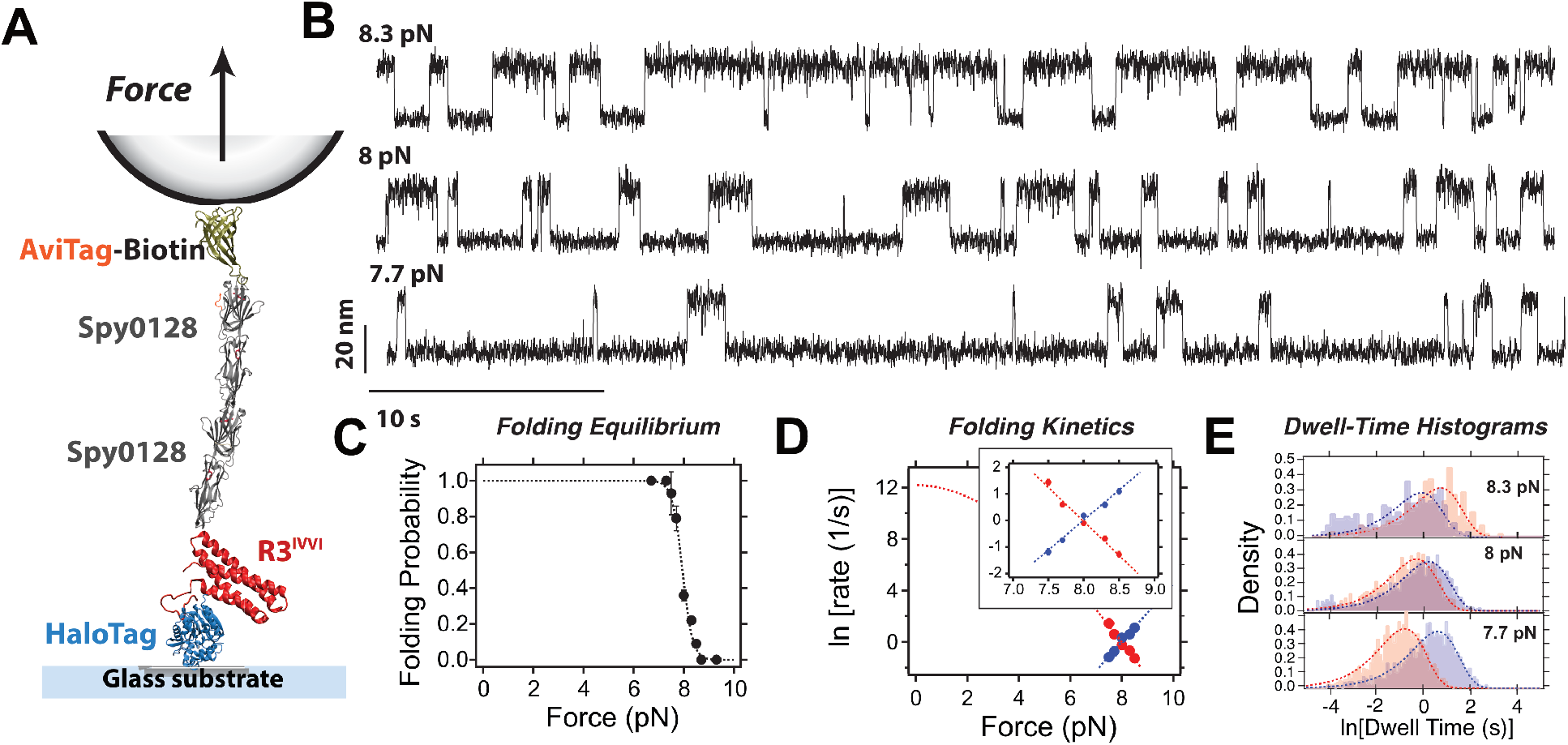
Complete nanomechanical mapping of R3^IVVI^: (A) Schematics of the magnetic tweezers and anchoring strategy for R3^IVVI^. (B) Fragments of the measured trajectory at three different forces, 8.3 pN, 8 pN, and 7.7 pN. R3^IVVI^ transitions in equilibrium between the folded (low extension) and unfolded (high extension) over a narrow force range, giving rise to changes on its extension of ~ 20 nm. (C) Folding probability of R3^IVVI^. The folding probability drops with force in a sigmoidal fashion over a force range of ~1.5 pN. Fitting parameters *F*_1/2_ = 7.95 ± 0.03 pN, *w* = 0.20 ± 0.02 pN. (D) Folding (red) and unfolding (blue) rates of R3^IVVI^ as a function of force. The scale employed highlights the narrow range that the rates span. (Inset) Detail of the folding rates over the 7-9 pN range. The folding rates follow a nearly exponential force dependence given the narrow force range that is explored. Fitting to Bell model yields 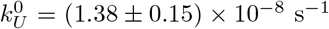 and 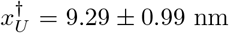 (unfolding) and 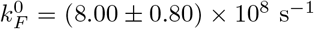 and 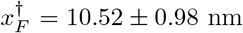. (E) Dwell-time histograms calculated with a logarithmic binning for 8.3 pN, 8 pN, and 7.7 pN (red, folded state; blue, unfolded state). We observe a single-exponential dependence at every force.

We first characterize the force response of R3^IVVI^ folding equilibrium and kinetics. Using 20 min-long JSON pulses between 7.3 pN and 8.7 pN, we calculate the folding probability (Fig. 3C) and the folding/unfolding rates (Fig. 3D). The folding probability drops steeply in a sigmoidal fashion, with a coexistence force of *F*_1/2_=7.95 ± 0.03 pN (*F*_1/2_, force at which the folded and unfolded states are equally populated). Due to this sharp force response, the folding/unfolding rates have a nearly-exponential dependence with force (Fig. 3D, inset), with very similar slopes. While the force dependence of the unfolding rates of a protein is usually well described by the Bell model,^29^ the folding rates *r*_F_ must account for the entropic elasticity of the unfolded state, which dominates the refolding transition.^24,30,31^ The force-dependence of the folding rates can thus be described by integrating over an energy profile dominated by the freely-jointed chain (FJC) model, leading to:^24,32^

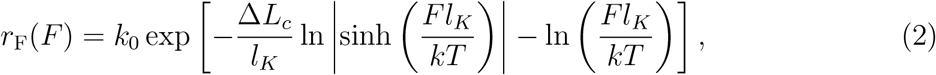

where is *k*_0_ the unfolding rate extrapolated to zero-force, Δ*L*_c_ the increment in contour length, and *l*_K_ the Kuhn length. In the case of R3^IVVI^, due to the narrow range of forces over which it folds/unfolds in equilibrium, the elastic contribution is nearly negligible, as seen in Fig. 3D, and the simple exponential Bell theory is a good approximation.

We continue our recording by studying in detail the folding dynamics at three different forces, 7.7 pN, 8 pN, and 8.3 pN, measuring six-hours-long pulses at each force. Figure 3E shows the dwell-time histograms in the folded (red) and unfolded (blue) states calculated with a logarithmic binning. Under this representation, standard in ion-channel analysis, a single-exponential distribution is transformed into a peaked distribution as *p*(*x*) = exp [*x* − *x*_0_ − exp (*x* − *x*_0_)], where *x* = ln (*t*), and *x*_0_ = ln (*τ*), being *τ* the average (folding or unfolding) time, which is the peak of the distribution.^19,33^ This transformation is useful to systematically identify multiple underlying kinetic processes, which can be hidden when representing the dwell-time distributions with linear binning. In our case, the three histograms fit a single exponential distribution, which indicates that the observed dynamics correspond to a simple Poisson process governed by a single folding/unfolding rate.

Our instrumental design balances the ability to measure single proteins for extended times and sampling its dynamics at a high temporal resolution. This recording was measured at 1352 Hz, which is sufficient to capture how the protein diffuses along its free energy profile and reconstruct this landscape by resolving low occupation regions utilizing thousands of transitions sampled at a high rate. Figure 4A shows a detail of the folding transitions of the measured R3^IVVI^. While a protein in equilibrium spends the majority of its time dwelling over the folded and unfolded basins, the brief excursions to switch between these states contain the most relevant information about a protein’s free energy landscape—dubbed the transition paths.^34,35^ We extract the transition paths for the folding and unfolding reactions at each force by identifying the fragments of trajectory where the protein escapes from the folded/unfolded basin (defined by their extensions, x_F_ or x_U_) to directly reach the unfolded/folded one (Fig. 4A, right). We measure the duration of the transition paths at each force (8.3 pN, 8 pN, and 7.7 pN) and calculate the histograms of transit times (Fig. 4B) for the folding (red) and unfolding (blue) transitions. At each force, the folding and unfolding transit time distributions are equal within error, as expected from the time-reversal symmetry.^36^ Unlike the folding and unfolding rates, which depend exponentially on the free energy barrier height Δ*G*^†^ and, hence, have a strong force-dependence, the transit times are dominated by the diffusion coefficient *D*, and the effect of the force is negligible. Assuming a harmonic barrier in the high-barrier limit, the average transition path time *τ*_TP_ can be expressed as 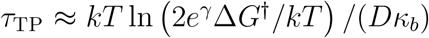, where *κ_b_* the stifness of the barrier top (assuming a harmonic barrier), and *γ* ≈ 0.577… the Euler-Mascheroni constant.^34^ As seen in Fig. 4B, we do not observe any appreciable change on the transition path time histograms with force, being the average transit time at every force *τ*_TP_ ≈ 11 ms. Following this approximation, the transition path time distribution *P*_TP_(*t*) decays exponentially as 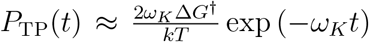 where 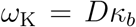.^34,35^ Both expressions provide a robust method to estimate the diffusion coefficient, given that the properties of the free energy landscape are known.

**Figure 4:**
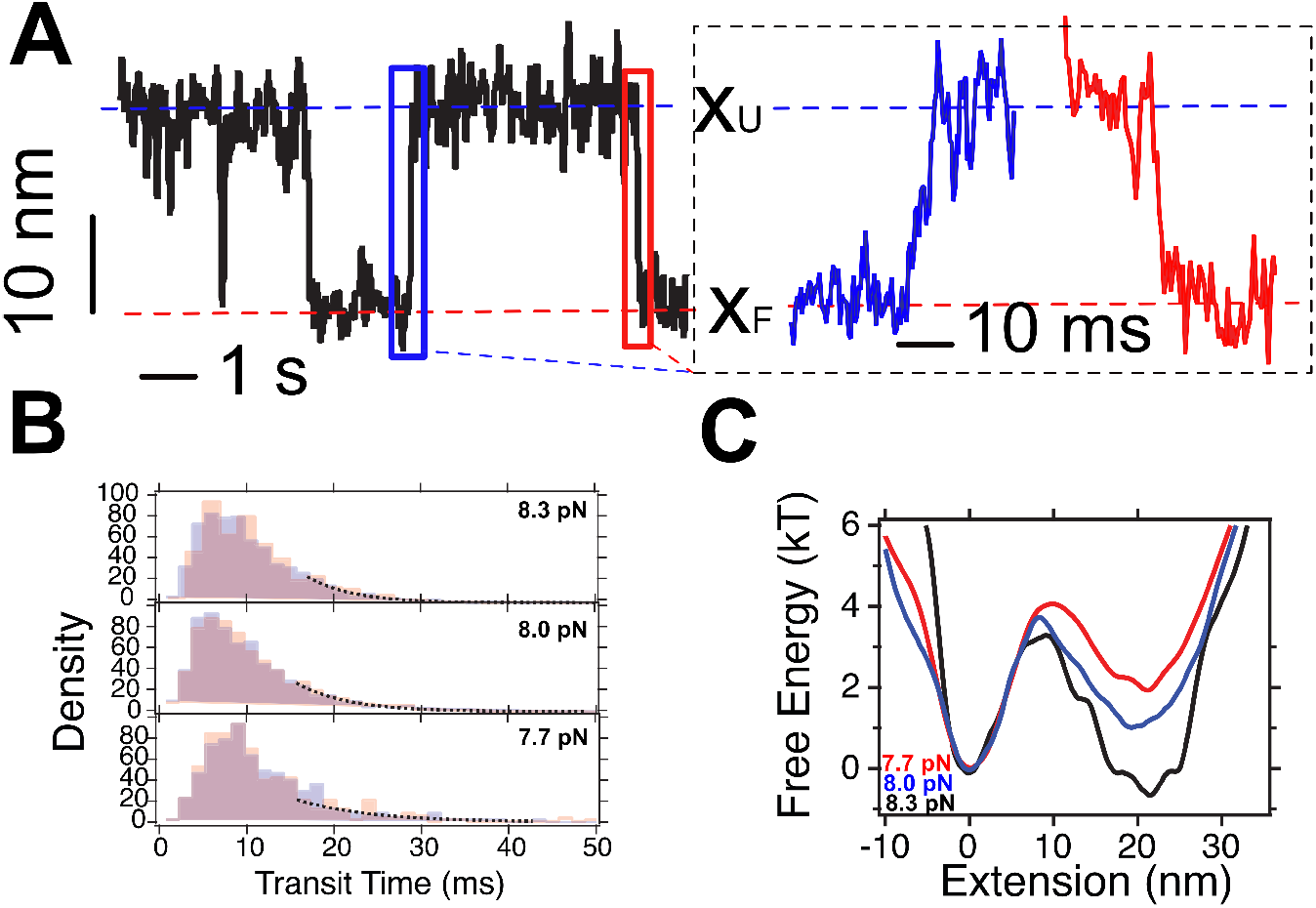
Transition paths and free energy landscape reconstruction: (A) Detail of R3^IVVI^ unfolding (blue) and folding (red) transition, with the unfolding (blue) and folding (red) transition paths highlighted. (B) Transition path time histograms at 8.3 pN, 8 pN, and 7.7 pN. The folded (red) and unfolded (blue) transit time histograms overlap, as expected from time reversal symmetry. The transition path times are insensitive to the magnitude of the pulling force (*τ*_TP_ ≈ 11 ms), over the explored range. (C) Free energy landscape of R3^IVVI^ at 7.7 pN (red), 8 pN (blue) and 8.3 pN (black).

To this aim, we reconstruct the free energy landscape *G*(*x*) at 7.7 pN, 8 pN, and 8.3 pN from the distribution probability of the molecular extension *p*(*x*) and its time derivative 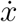 as:

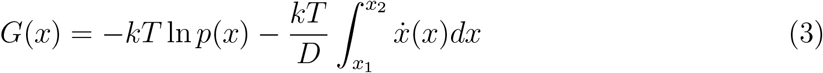

being *x*_1_ and *x*_2_ the range of *x*. This equation accounts for the equilibrium contribution (first term, Boltzmann inversion) and the non-equilibrium contribution (second term).^37,38^ While this latter term is crucial when employing non-equilibrium trajectories, in our case, given the symmetry between the folding and unfolding transition paths, it is negligible. In practice, as it has been extensively discussed,^39,40^ the measured dynamics do not correspond to the real molecular dynamics, but include the convolved instrumental influence, so the direct application of Eq. (3) on the measured signal does not result in the actual molecular free energy landscape. In such a case, the molecular free energy landscape is calculated by estimating an instrumental point-spread-function and using a deconvolution algorithm to remove the instrumental effect.^39^ Here, and unlike the case of optical tweezers and AFM, we do not have a direct effect of the force probe due to the negligible stiffness of the magnetic trap (≪ 10^−6^ pN/nm) that results in the hallmark force-clamp conditions of magnetic tweezers. However, the image analysis algorithm we employ to infer the molecular extension introduces instrumental noise to our recordings that smears the actual protein dynamics. To deconvolve this effect, we estimate the point-spread-function of our experiment by measuring long recordings of two non-magnetic beads, which are tightly attached to the glass surface and do not respond to force, hence, mimicking the noise introduced by our image analysis algorithm. As shown in Fig. S6, the point-spread-function is a Gaussian distribution with a standard deviation of 1.7 nm.

Figure 4C shows the free energy profiles at 7.7 pN, 8 pN, and 8.3 pN calculated with the described procedure. Force tilts the free energy landscapes towards the unfolded state, hence, stabilizing this conformation. Due to the small range of forces explored by R3^IVVI^, we only observe a small increase in the extension change upon unfolding (19.5 nm at 7.7 pN, and 20.3 nm at 8.3 pN). However, we readily observe the differences in the width of the unfolded basin that arise from the gain in contour length, leading to a very different stiffness, *κ_F_* = 0.28 ± 0.02 pN/nm (folded) and *κ_U_* = 0.08 ± 0.01 pN/nm (unfolded) (at 8 pN, see SI Appendix for the free energy landscapes properties at all forces). In a simple two-state scheme, to a first approximation, the force is assumed to act linearly with a -*F* · *x* term. The landscapes estimated for R3^IVVI^ are in agreement with this approximation, as subtracting the landscapes at 8.3 pN and 7.7 pN recovers this linear contribution, which has a slope of ~ 0.6 pN, the difference in force between these two landscapes (Fig. S7).

From the properties of the free energy landscapes, we can estimate the diffusion coefficient of the measured dynamics in two different ways. First, from the average transition time and since the stifness of the free energy barrier is *κ_b_* = 0.37 ± 0.04 pN/nm and the barrier height 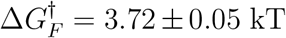 (using the data at 8 pN), we obtain a first estimation of the diffusion coefficient *D* ≈ 2870 ± 690 nm^2^/s. Second, by fitting the tail of the transition path time distributions (knowing 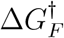), we obtain that the fundamental frequency is *ω_K_* = 273.3 ± 4.2 s^−1^, which leads to *D* ≈ 3032 ± 331 nm^2^/s. In this sense, both methods provide very similar estimates for the diffusion coefficient. With this value of the diffusion coefficient and the properties of the free energy landscape, we can estimate the folding and unfolding rates using Kramers’ reaction rate,^41^ which yields *r_U_* ≈ 0.86 s^−1^ and *r_F_* ≈ 1.28 s^−1^, in excellent agreement with the measured rates (Fig. 3D), which supports the consistency of the estimated free energy landscapes and values of the diffusion coefficient. Notably, the diffusion coefficient of a superparamagnetic bead is over two orders of magnitude larger than the diffusion coefficient of R3^IVVI^ dynamics, which suggest that the protein folding dynamics are rate-limiting, as previously suggested.^22^

### Nanomechanics of protein L

Protein L—a superantigen bacterial domain with a topology that comprises four *β* strands packed against an *α* helix—is a common protein folding model, and has been used as a molecular template to calibrate single-molecule instrumentation.^16,17,19^ Protein L has slow folding dynamics between forces of 4 and 10 pN, which arise from the hysteresis of its folding and unfolding forces—unfolding quickly at forces >40 pN, while collapsing and folding fast below 4 pN.^42^ Owing to the slow kinetics of protein L folding dynamics, its nanomechanics have been mostly characterized using chimeric polyprotein constructs that amplify the signal. To characterize in detail the folding dynamics of a single protein L individual, we measure one protein L monomer for five uninterrupted days (Fig. 5A). Our full recording comprises initial short pulses at high force to measure its unfolding kinetics and long JSON-pulses (~8 hours) at low force to characterize its folding properties. Figure 5B shows short fragments of protein L dynamics at 7.5 pN, 6 pN, and 4.5 pN, which highlights the much slower folding kinetics compared to R3^IVVI^. Since protein L exhibits reversible folding dynamics over a wide force range, we can readily observe the impact of polymer elasticity on its folding mechanism as the force scaling of its extension changes (step sizes), being ~5 nm at 4.5 pN, while ~8.5 nm at 7.5 pN.

**Figure 5:**
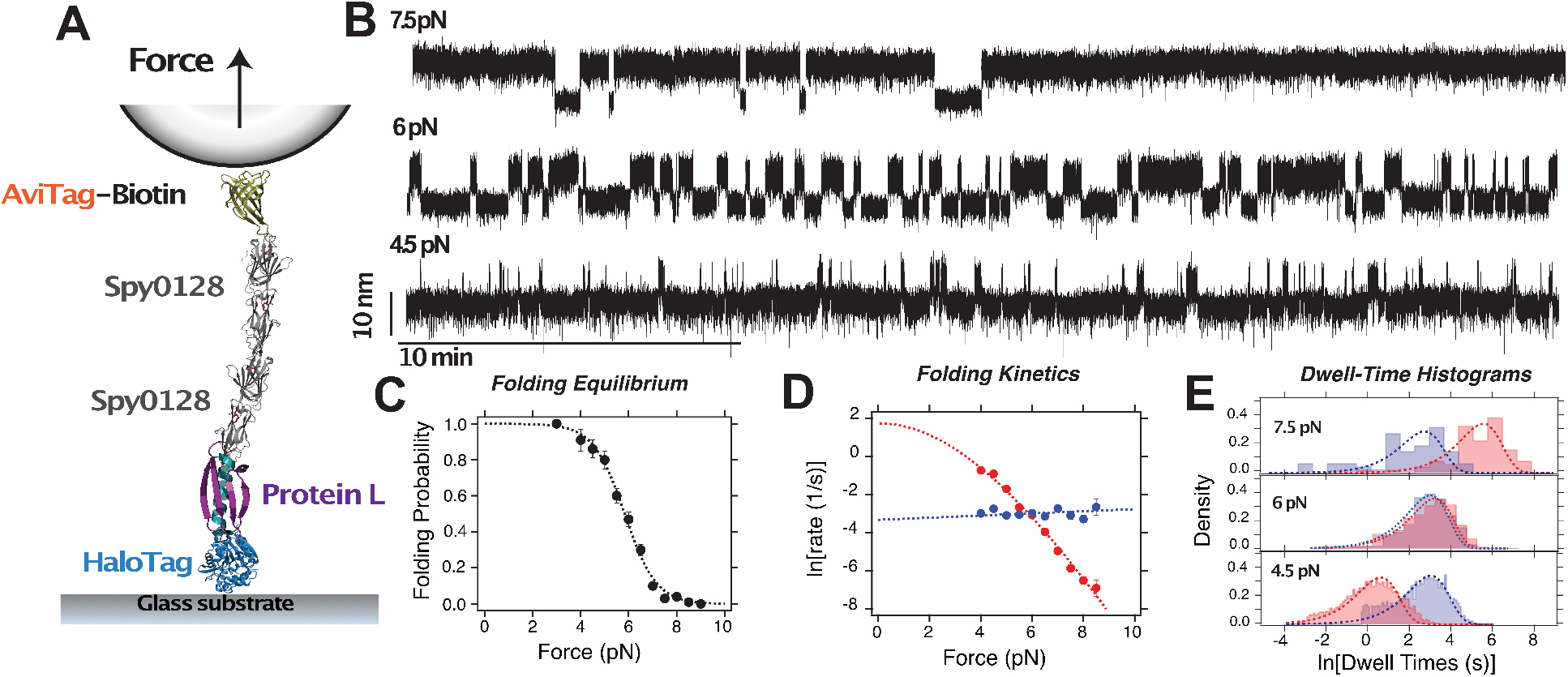
Complete nanomechanical mapping of a single protein L domain: (A) Schematics of the magnetic tweezers and anchoring strategy for our protein L experiments, identical to the one employed for R3^IVVI^. (B) Fragments of the measured trajectory at three different forces, 7.5 pN, 6 pN, and 4.5 pN. In contrast to R3^IVVI^, protein L folding dynamics explore a much wider range of forces with slower kinetics, that lead to completely different nanomechanical properties. (C) Folding probability of the protein L molecule. The folding probability drops with force following a sigmoidal dependence, showing reversible folding dynamics between 4 pN and 9 pN, approximately. Fitting parameters *F*_1/2_ = 5.83 ± 0.04 pN, *w* = 0.65 ± 0.04 pN (D) Folding (red) and unfolding (blue) rates of the protein L molecule as a function of force. The force dependency of the rates is strongly asymmetric; the folding rates drop quickly with force, while the unfolding rates have a nearly flat force-dependency. Additionally, the folding rates show a non-exponential behavior due to the contribution of the elasticity of the unfolded state on the refolding transition. Fitting the folding rates to Eq. (2) leads to 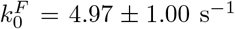, *L_c_* = 14.82±6.4 nm, and *l_K_* = 0.97 ± 0.52 nm, these latter two parameters in great agreement with the known FJC properties of protein L. Fitting the unfolding rates to the Bell model yields 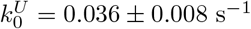 and *x*^†^ = 0.23 ± 0.02 nm. (E) Dwell-time histograms calculated with a logarithmic binning for 7.5 pN, 6 pN, and 4.5 pN (red, folded state; blue, unfolded state). We observe a single-exponential dependence at every force.

Similar to the scheme developed in Fig. 3, we characterize first the force-response of protein L through the folding probability (Fig. 5C) and the folding/unfolding rates (Fig. 5D). The folding probability also has a sigmoidal dependence, but with a shallower force dependence, showing a coexistence force of *F*_1/2_=5.83 ± 0.04 pN. The difference between the nanomechanics of R3^IVVI^ and protein L is distinctly reflected in the force-dependency of its folding and unfolding rates that, as shown in Fig. 5D, are highly asymmetric. The unfolding rates (blue) are nearly flat over the folding range, which, accounting for its rates measured at high forces (not shown), leads to a distance to the transition state of *x*^†^ = 0.23 ± 0.02 nm, assuming Bell model. By contrast, the folding rates (red) drop very quickly with force, following a non-exponential shape as predicted by Eq. (2), readily suggesting that the folding transition is dominated by the entropic elasticity of the extended polypeptide chain. The dwell-time histograms (Fig. 5E) are exponentially distributed, with the dwell-time in the folded state (blue) remaining nearly unchanged, while the time spent in the unfolded state (red) increases very quickly with force.

Figure 6 shows a detail of protein L folding dynamics. Although these dynamics appear similar to those observed for R3^IVVI^, the folding mechanism of protein L strongly differs. To unveil this process, we extract the transition paths, and calculate the transition path time histograms (Fig. 6B), which also fulfil the time-reversal conditions and maintain an apparent insensitivity with force. However, the average transition times increase considerably for protein L (*τ*_TP_ ≈ 33 ms) compared to R3^IVVI^, which could suggest slower diffusive dynamics.

**Figure 6:**
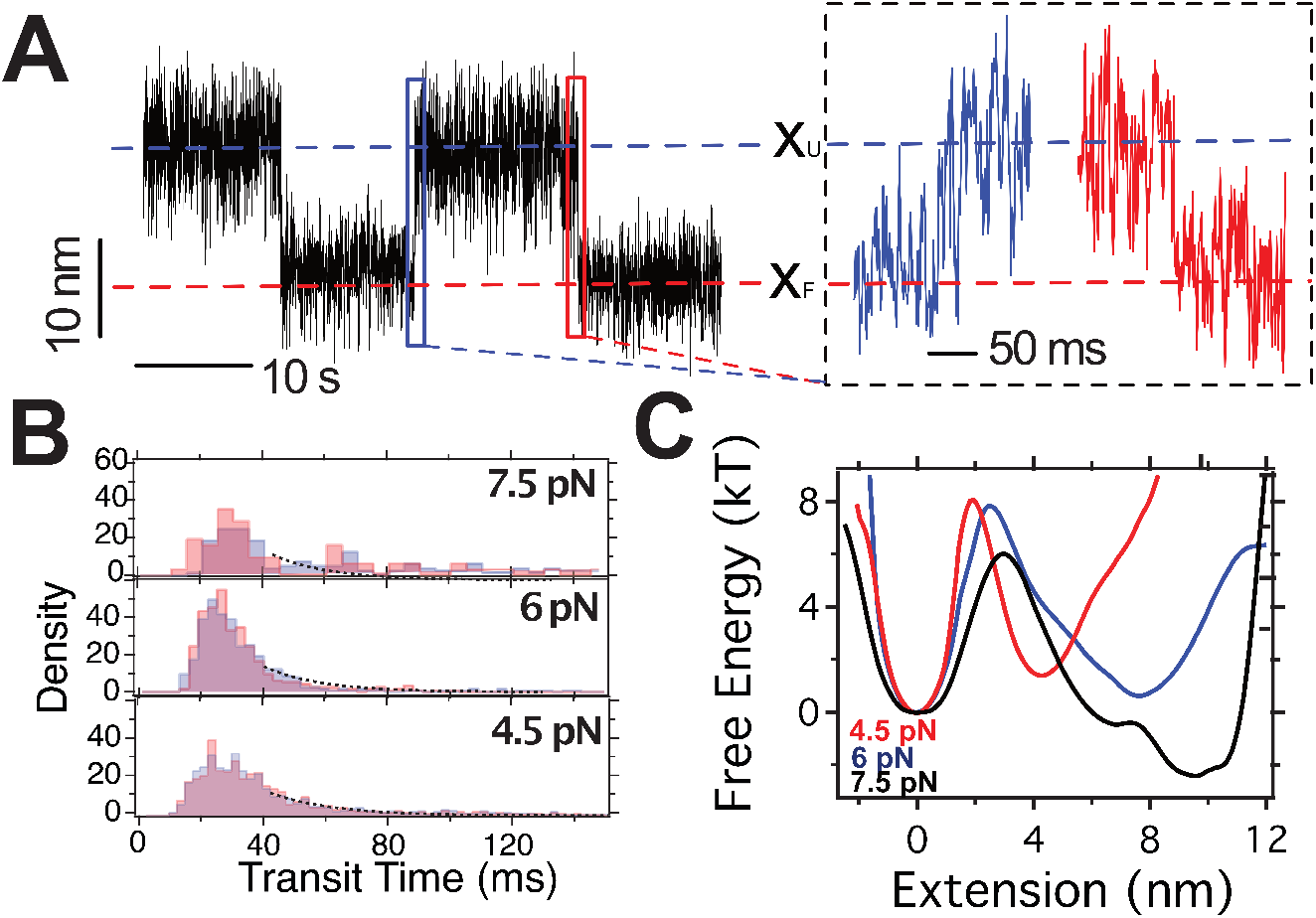
Transition path times and free energy reconstruction of protein L: (A) Detail of protein L unfolding (blue) and folding (red) transitions. (B) Transition path time histograms at 7.5 pN, 6 pN, and 4.5 pN. The folded (red) and unfolded (blue) histograms overlap, as expected from time reversal symmetry. The average transition time is nearly 3 times higher than those of R3^IVVI^ (*τ_TP_* ≈ 33 ms). (C) Free energy landscapes at 4.5 pN (red), 6 pN (blue) and 7.5 pN (black). The shape of the free energy landscapes and how they are modulated by force illustrate that the folding mechanics of protein L are not well-understood with a simple two-state model. The unfolded state position shifts with force following the non-linear FJC model, and also widening significantly with force. This indicates that the effect of the force on the folding landscape cannot be ascribed to a simple linear tilt.

Following the deconvolution procedure, we reconstruct the free energy landscape at the measured forces. Figure 6C shows the free energy landscapes along the molecular extension at 4.5 pN (red), 6 pN (blue) and 7.5 pN (black), which manifest the great differences between R3^IVVI^ and protein L folding under force. First, the reconstructed free energy landscape captures both the asymmetry of the landscape and the notable shift in the position of the unfolded state, which scales non-linearly following the FJC, with ~5 nm (4.5 pN) and ~9 nm (7.5 pN). In this regard, it is clear that a linear approximation does not account for the actual effect of the force on the free energy landscape, and subtracting the landscapes leads to non-linear contributions that do not resemble the classical linear tilt (SI Appendix, Fig. S7). When estimating the diffusion coefficient *D*, we obtain surprisingly low values of *D* ≈ 71.9 ± 4.7 nm^2^/s (from *τ*_TP_), and *D* ≈ 67.4 ± 3.5 nm^2^/s (from the fits to *P*_TP_). Using these estimations, the Kramer’s rate at 6 pN is *r* ≈ 0.0057 s^−1^, while the measured ones are nearly 10 times higher. This discrepancy arises likely due to protein L folding mechanism, which is dominated by the entropic elasticity of the unfolded state, eventually leading to the failure of theories developed for simple analytic two-state landscapes.

### Characterizing single-protein nanomechanical diversity

While single-molecule approaches enable to interrogate individual biomolecules, it is typically necessary to assess several molecular specimens in order to accumulate enough data for a robust statistical significance. This approach is intrinsically based on the ergodic hypothesis, which states that the time average of a physical observable equals its ensemble average.^43^ Here, we have undertaken this first approach, and thoroughly mapped the nanomechanics of two different proteins by measuring their dynamics for long timescales and time-averaging their properties. However, an inherent assumption of the ergodic hypothesis is that all the individual components of the ensemble are identical entities so that all trajectories employed for the ensemble average transverse the same phase space. Here, we can directly test this hypothesis and compare the nanomechanical performance of different single molecules to look into their functional heterogeneity.

We measure several individuals of R3^IVVI^ and protein L and compare their response to force. Figures 7A and B show the folding dynamics of R3^IVVI^ (A) and protein L (B) for two different protein specimens subject to the same pulling force. Interestingly, each protein individual shows a remarkably different behavior at the same force, populating differently the folded/unfolded states and exhibiting different transition kinetics. Hence, the comparison of these recordings readily suggests a degree of dispersion in the nanomechanical properties of each protein L and R3^IVVI^ molecule.

**Figure 7:**
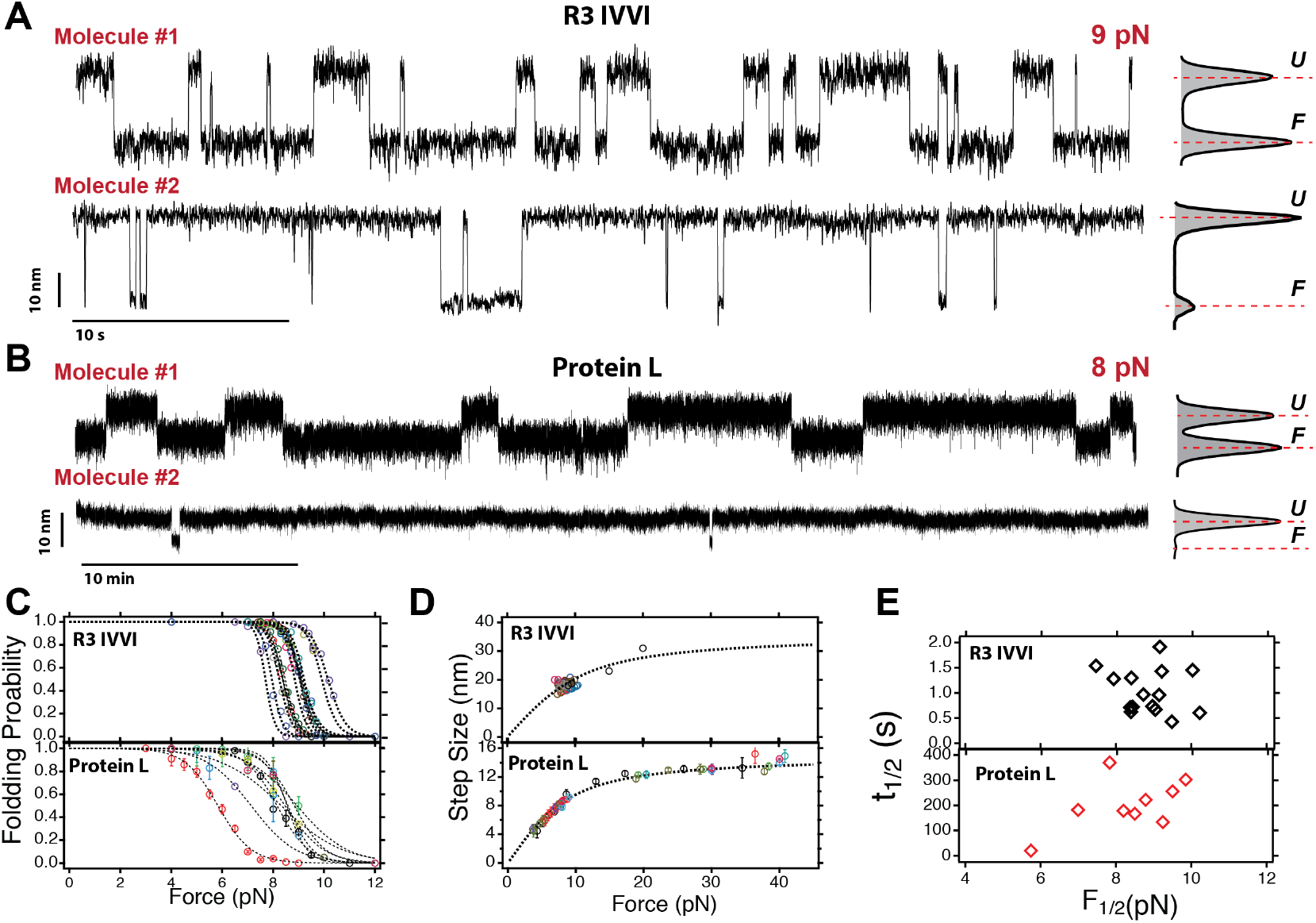
Characterization of the nanomechanical identity among protein individuals: (A) Fragment of R3^IVVI^ folding mechanics close to *F*_1/2_ measured for two different R3^IVVI^ molecules. (B) Fragment of protein L folding mechanics close to *F*_1/2_ measured for two different protein L molecules. (C) Folding probabilities for R3^IVVI^ (upper) and protein L (lower) measured over different protein individuals (each color/symbol is a different molecule). We observe in both cases a dispersion in the folding probability force dependency, which is more evident in the case of protein L. (D) Step-sizes as a function of the pulling force for R3^IVVI^ (upper) and protein L (lower). While it is only possible to measure the step-sizes of R3^IVVI^ over a narrow range of forces, all the protein L molecules have step-sizes that fall on a master FJC curve, which indicates that our setup applies well-calibrated and reproducible forces, which do not explain the dispersion in the nanomechanical properties we observe. (E) Scatter plot of the coexistence force *F*_1/2_ and average coexistence dwell-times *t*_1/2_ for R3^IVVI^ (black) and protein L (red).

To further characterize this observation, we measure the folding probability of several R3^IVVI^ (16 molecules) and protein L (9 molecules) (Fig. 7C), which readily shows the heterogeneity in the mechanics of each protein individual. In the case of R3^IVVI^, the folding probabilities have a sigmoidal dependence with *F*_1/2_ that spans between ~ 8 pN and ~10 pN, while for protein L the dispersion in *F*_1/2_ spans between ~6 pN and ~10 pN. Notably, the differences in the folding probabilities do not correspond to simple shifts of the sigmoidal curve, but their shape also differs amongst them, showing a different force-dependency over the full folding range. To demonstrate that these differences do not arise from a poor calibration of our setup or from the size-dispersion in the magnetic beads (M-270 beads have a size dispersion of ~2 %), we measure the step sizes of R3^IVVI^ and protein L. This molecular observable depends on the contour length of the molecules, this is, the number of residues in the protein sequence, and, as previously demonstrated, is an extremely force-sensitive metric that can be employed as a calibration strategy.^17,19^ Figure 7D shows the step sizes for R3^IVVI^ (upper) and protein L (lower) as a function of force. Unfortunately, R3^IVVI^ explores a minimal range of forces, and its step sizes can only be ascertained over a narrow force range, since R3^IVVI^ unfolding becomes too fast at forces above 15-20 pN to resolve the event. By contrast, protein L, owing to its slow unfolding kinetics, allows us to measure its step sizes between 4 and 40 pN, and fully account for their force dependency. In contrast to the folding probability, the step sizes of every protein L molecule fall on the same curve given by the FJC model with the previously determined properties, Δ*L_c_* = 16.3 nm and *l_K_* = 1.1 nm, readily indicating that our force calibration and bead-to-bead variations allow us to apply forces in an accurate and reproducible way and that the observed dispersion is an intrinsic property of the measured molecules.

To portray the nanomechanical heterogeneity among different R3^IVVI^ and protein L individuals, we measure their folding/unfolding kinetics and determine their coexistence force *F*_1/2_ and their dwell times at this force *t*_1/2_. To accurately estimate these values, we obtain them from the intersection point between the folding/unfolding rates by fitting their force-dependency (SI Appendix, Figs. S8 and S9). Figure 7E shows a scatter plot of *F*_1/2_ and *t*_1/2_ that illustrates the heterogeneity on the folding mechanics of each measured protein. In both cases, we observe a significant dispersion in the values of *F*_1/2_ and *t*_1/2_, more pronounced in the case of protein L. R3^IVVI^ molecules have an average coexistence force of ⟨*F*_1/2_⟩ ≈ 8.7 pN and ⟨*t*_1/2_⟩ ≈ 1.3 s, both values in agreement with previous work,^44^ while the dispersion of these averages (standard-deviations) is *δF*_1/2_ ≈ 1 pN, *δt*_1/2_ ≈ 0.75 s. For protein L, we measure ⟨*F*_1/2_⟩ ≈ 8 pN, and ⟨*t*_1/2_⟩ ≈ 200 s, with a dispersion of *δF*_1/2_ ≈ 2.4 pN, *δt*_1/2_ ≈ 150 s. Interestingly, the average properties of protein L are in agreement with measurements of the protein L octamer, which could suggest that the use of polyproteins offers a “pre-averaged” signal, that reduces the apparent nanomechanical heterogenity.^42^ Overall, while the step sizes, an intrinsic property of the polypeptide chain, are homogeneous among different molecules, the properties related to their folding landscape show a high degree of heterogeneity.

## Discussion

The inception of single-molecule techniques opened the gates to reconstructing distributions of molecular properties, and appreciate the dynamic and heterogeneous nature of biomolecular behaviors. These advances offered valuable insight on the functional heterogeneity in diverse biomolecular systems, such as enzymatic reactions, ligand binding, or molecular motors.^45–47^ Force-clamp atomic force microscopy (AFM) experiments evidenced the existence of protein heterogeneity in poly-ubiquitin molecules, which exhibited non-exponential unfolding kinetics likely due to a distribution of free energy barrier heights—a manifestation of the so-called static disorder in the Zwanzig’s terminology.^5^ More recently, a systematic study of several single-molecule force spectroscopy data-sets suggested a widespread mechanical heterogeneity in different protein, nucleic-acid, and complex-ligand systems.^48^ While these works offered indirect evidence of functional heterogeneity among single proteins, instrumental limitations have prevented to directly address this question, which would require a direct comparison between the properties obtained from time averages over long trajectories and those that arise from ensemble averages over several molecules; this is, to test if the ergodic hypothesis holds in protein nanomechanics.^43^

Our instrumental development opens up the gates to profile the nanomechanics of an individual protein by collecting extensive recordings of its dynamics and accumulating enough events to fully portray its molecular properties. Due to this capability, we have directly captured the mechanical diversity of two protein domains, the talin R3^IVVI^ domain and protein L. This heterogeneity is manifested as a dispersion of their coexistence forces *F*_1/2_ and their folding/unfolding kinetics. While the potential impact of such protein diversity in a physiological setting is unclear, it is remarkable to note how the functional heterogeneity of RNA molecules has been widely reported, while this observation on single proteins has remained far more elusive.^49–51^ Hence, it seems evident that the diversity of RNA phenotypes is far more obvious, compared to that of proteins. It is enticing to think that RNA is a more “immature” and ephemeral biomolecule (from the evolutionary perspective), while proteins have an optimized structure and their synthesis undergoes further proofreading process, which could narrow the distribution of functional molecules. However, a protein’s response to a stretching force is a key determinant of many biological processes. For example, in the case of talin, its folding status regulates the binding of ligand partners such as vinculin.^44,52^ More generally, the mechanical stability of proteins has been suggested to tune its import rate through the nuclear pore complex, the gateway of nuclear translocation.^53^ Hence, it could be argued that uniform mechanical properties are a desirable molecular property, specially in the case of mechanosensing proteins. However, it remains an open question whether the observed mechanical diversity also applies to proteins operating in the cell, or biological control processes can further filter protein populations by its mechanical properties.

From the physical perspective, the exact origin for such nanomechanical dispersion is uncertain. As some theoretical and computational protein folding models have explored, biomolecular free energy landscapes can be highly rugged, containing a degenerate ground state with multiple functional minima separated by large barriers.^5,48,54^ In such a case, while bulk techniques would measure the average conformation of such ensemble of states, singlemolecule approaches would sample independently biomolecules “frozen” at each individual state. However, the observed dynamics are robust and each individual protein maintains, within statistical significance, a “nanomechanical” identity. This would imply that, if the rugged landscape model is correct, such multiple states are separated by extremely high barriers, inaccessible even over timescales of several days. Additionally, a protein is a dynamic and evolving molecule, which is likely to undergo multiple and random chemical modifications that can alter in a more or less significant way its free energy landscape. As we previously demonstrated, mechanical stretching exposes cryptic residues that can be irreversibly oxidized, which eventually blocks protein folding and leaves an unstructured and labile polypeptide.^42,55^ By contrast, these random chemical modifications on residues natively exposed to the solvent are less detrimental on the protein’s fold, while, however, could still render a modified, yet functional, landscape.^56^ Importantly, our data clearly indicates that the length of the protein sequence is a constant among the different molecular specimens, as demonstrated by the high homogeneity of the polymer elasticity properties of the unfolding protein L domains, which also verifies our ability to apply calibrated forces and the unambiguous fingerprint for our tethers.

Our data further demonstrates the key role of polymer physics on protein folding under force. The entropic elasticity of the unfolded polypeptide dominates the refolding transition, owing to the mechanical work that protein folding must do in order to collapse against the pulling force.^24,30,57^ The most obvious manifestation of this physical property is the change in the step-sizes with force following standard polymer physics models such as the FJC.^58^ Another prediction from the FJC that our recordings readily show is the increase in the magnitude of the end-to-end length fluctuations upon unfolding, due to the gain in contour length that results in a wider free energy basin. This property is captured now in the reconstructed free energy landscapes (Figs. 4 and 6), where the stiffness of the unfolded state is reduced by up to 70% for R3^IVVI^ (contour length increment Δ*L*_c_ ~ 40 nm) and 55% for protein L (contour length increment Δ*L*_c_ ~ 16.3 nm). The recognition of the fundamental role of polymer physics on a protein’s folding landscape has often remained disregarded, likely since most of the studied proteins explore narrow force-ranges which make valid two-state-like theories based on linear approximations of the effect of force. In this sense, these properties are clearly reflected when comparing the nanomechanics of R3^IVVI^ and protein L. Similar to nucleic-acid hairpins or proteins with fast hopping kinetics, R3^IVVI^ shows reversible folding dynamics over a range of ~1 pN, which result in a negligible change on its step sizes. Its folding/unfolding rates have an exponential dependence following the Bell model and resulting in a good agreement of its properties with theories developed for simple bistable potentials, which indeed resemble its free energy landscape (Fig. 4). In this sense, R3^IVVI^ folding is likely driven by the hydrophobic collapse of the unfolded polypeptide, with minimal contribution of short-range bonds, resulting in a compliant free energy landscape. By contrast, protein L, akin to other proteins with Ig-like or *α/β* folds, has a very cooperative and brittle fold, as reflected by a very short distance to transition state for unfolding (<1 nm). While the unfolding rates increase very slowly with force, the refolding rates have a very steep and non-exponential dependence with force, which can be well explained assuming that this transition is dominated by the free energy surface dictated by the FJC, as our fit to Eq. (2) suggests. As our free energy reconstruction demonstrates, the free energy landscape of protein L is indeed highly asymmetric, which readily demonstrates the non-linear effect of force in the shift of the unfolded state position due to the contribution of stretching an unfolded polypeptide. As a consequence of this contribution, the theories based on two-state-like descriptions of protein folding fail to explain protein L folding dynamics. In this sense, thanks to the compromise between high-resolution and stability of our magnetic tweezers setup, we have been able to reconstruct the nanomechanics of two different proteins and directly demonstrate the inarguable contribution of polymer physics in protein folding under force.

In summary, we have here introduced a novel magnetic tweezers development, which combines a simple and affordable design with improved resolution and stability that allows the full characterization of individual proteins. Our approach is grounded on our previous developments, implementing the use of a tape head for enhanced force control and stability,^19^ and the use of HaloTag chemistry to achieve highly specific and long-term tethers.^17^ The introduction of a high-precision mounting piece for the tape head results in an enhanced and reliable control of the pulling force that involves no moving parts, which further increases the instrumental stability. Furthermore, the simplified and dedicated microscope design, without relying on standard multi-functional microscopes, results in an increased signal-to-noise ratio and negligible drift. These properties combined enable the acquisition of long recordings that accumulate a high number of molecular events, and resolving at the same time the nuances of the free energy landscape. Importantly, our setup highlights its bespoke and affordable design, which facilitates its assembly and installation following the details and instructions of the SI Appendix. Hence, we anticipate that our magnetic tweezers approach has the potential to become a widespread tool to study biomolecules under force, and provide insights on the relevance of mechanical regulation in biological processes including cellular adhesion, mechanotransduction, or tissue mechanics.

## Acknowledgement

We thank David Scott for his help with the 3D-model of the setup (Fig. 1). This work was supported by NIH Grant R35129962. R.T-R. and A.A-C. acknowledge Fundacion Ramon Areces for financial support.

