## Supplementary material for "Identical Sequences, Different Behaviors: Protein Diversity Captured at the Single-Molecule Level": SI Appendix

Rafael Tapia-Rojo,[1,*] Alvaro Alonso-Caballero,[1] Carmen L. Badilla,[1] and Julio M. Fernandez[1]
[1]*Department of Biological Sciences, Columbia University, New York, NY 10027, USA*

### MAGNETIC TWEEZERS; SETUP DETAILS

Our magnetic tweezers design (Fig. 1 main text) is assembled from off-the-shelf and custom-made components for an affordable cost. This instrument is based on our previous designs that included a regular inverted microscope. Here, we design a simple dedicated microscope that greatly simplifies the setup and reduces its cost. We include here a guide of assembly, including details about the microscope design, the magnetic tape head implementation, and the electronics.

#### Microscope design

For the optics of our setup, we use a 160 mm objective mounted on a piezo electric actuator (element #4, P-725, PI), which is on top of the Z-stage (element #5, Thorlabs). While the Z-stage can be used to do coarse movements of the objective (for example, after mounting the fluid chamber to find the focal plane), the piezo electric actuator is used to focus at high-precision, allowing to track the position of the magnetic bead with nm resolution. Figure 1 shows a diagram for the optical pathway of our microscope design, separating the illumination pathway (yellow) and the image (orange). The colors of the beams are arbitrary. The sample is illuminated from below using a white LED that provides a point light source (element #12, Thorlabs). The light beam is then collimated using a plano-convex lens (element # 10, Thorlabs) and then focused on the 160 mm objective (element #3, Zeiss) using a tube lens (element #7, Thorlabs), after passing through a green filter (element #9, Thorlabs) and an iris diaphragm (element #8, Thorlabs). The filter helps in producing sharper interference rings, while removes high frequencies that increase the production of reactive oxygen species in the protein solution, while the iris modulates the aperture of the microscope. The virtual image created by the objective is directed to the CMOS camera using a 50:50 beam splitter (element #6, Thorlabs), and converted to a parallel image using a plano-convex lens (element #15).

The extension of the protein is tracked in real time using the image analysis algorithm previously described [1]. Briefly, at the start of the experiment, we build a stack library of the radial vectors of the chosen magnetic and reference beads with a 2,560 $\mu$m travel with 20 nm steps (128 radial vectors) using the piezo actuator. The radial vector is calculated from the Fourier transform of images of beads, by summing over the intensity of the pixels at equal radial distance (removing low frequencies to account only for the beads' rings). During the course of an experiment, the real-time radial vector of the magnetic and reference beads is compared to the stack library by calculating the Pearson cross-correlation. To obtain the change in extension of the protein (height of the magnetic bead compared to the reference one), we fit a Gaussian to the correlation vectors using the analytical Caruana's algorithm [2], and infer the extension from the difference in position of the peaks as obtained from the fit. Every 1,000 frames the drift is corrected by refocusing the objective to keep the reference bead on focus following its radial vector at the start of the experiment.

#### Implementation of the magnetic tape head

Our magnetic tweezers design implements a magnetic tape head (element #17), which allows generating pN-level forces with a resolution and control superior to other implementations of electromagnets or permanent magnets. As previously described, the magnetic field generated by the tape head can be described analytically, which allows to develop a theory for the pulling force as a function of the distance from the head gap to the magnetic probe and the electric current supplied to the head [3]. In our novel magnetic tweezers setup, we maintain the tape head at a fixed distance, and control the force only through the electric current. The pulling force changes over a length scale of the gap size, 25 $\mu$m, which requires an accurate positioning in order to generate calibrated and reproducible forces. To this aim, we designed a mount piece that is fabricated with high-precision CNC (element #16), which maintains the tape head fixed 300 $\mu$m away from the magnetic probe, greatly increasing the reliability and stability of our setup, since no moving parts are required. Figure 2 shows a technical drawing indicating the dimensions of the piece (in mm), designed for the rail-carriages and tape head we employ (see Table I). The design includes the holes for mounting it on the optical rail carriages (element #2) and maintaining the tape head gap aligned with the optical axis of our microscope. The tape head is mounted with a dowel-pin system that fixes it position. In this assembly, the gap is exactly 450 $\mu$m away over the surface so that, when using 150 $\mu$m-thick bottom glasses (Ted Pella), the gap is 300

$\mu$m from the magnetic beads. When using the M-270 beads, the force law is [3]:

$$F(I) = (2.79 \times 10^{-5})I^2 + (1.64 \times 10^{-2})I, \tag{1}$$

where $I$ is the electric current in mA. The tape head saturates when using electric currents over 1 A. Hence, we can generate mechanical forces between 0 pN, and 44 pN. We have recently developed an implementation of the larger M-450 magnetic probes which pushes the upper force limit to ∼220 pN [4].

#### Electronic circuit

The tape head is maintained under electronic feedback, which enables to do swift force changes and apply complex force signals [3, 5]. We designed a custom PI circuit (proportional+integral) to maintain the electric current (force) under feedback (see Fig. 3 for its scheme). Since the tape head is an inductance impedance, we use a high-precision 2 Ω resistor (R17, Fig. 3) in series with the head and maintain the voltage drop across it under feedback, and, hence, the electric current through the head.

The circuit has two independent inputs (commands) from two separate DAQS, one accounting for the DC force (signals-3-4), and one for the AC component (signals-5-6). This implementation allows applying complex force signals such as oscillations or mechanical noise, as in [5]. The use of this second DAQ is optional, and the system can work only with a single command if no time-dependent force signals are to be applied. Both command signals are summed by a summing amplifier (A0) that feeds the error amplifier (A2). The other input is the voltage drop across the high-precision resistor (R17), which is also read as an output with a follower amplifier (A1). The error signal (difference between the command and the actual signal) goes to the PI circuit (proportional A3 and integral A4), calibrated for the used tape head. The signal has an output gain (A5) and is finally amplified by a power amplifier (LM3886), which supplies the desired current to the head. We use a pair of 12 V batteries as power supply, which provide a clean and fast source electric current, crucial for doing fast force changes. This implementation allows changing the force over ∼40 $\mu$s, only limited by the slew-rate of the power amplifier.

We include in Fig. 3 the board layout for surface mounting and a picture of the circuit once assembled.

#### Fluid chamber assembly and manipulation

Our fluid chambers (element #19) are custom fabricated using two glass cover slides (Ted Pella; 40x24 mm bottom, 22x22 top), which are functionalized as described below. The cover slides are sandwitched with a laser-cut parafilm pattern, and the assembly is melted with a hot plate with weight on top. The fluid chamber is clipped to a custom-design manipulation fork (element #18) fabricated by metal-stamp using two stripes of double-sticky tape. The fork is mounted on a XY stage (element #20) using two high-precision shoulder screws that allow horizontal and vertical displacement over a ∼3x3 mm region. The explored region can be changed by displacing horizontally the fork using its slit.

#### List-of-parts

We include in Table I a complete list of parts, following the numbering of Figure 1 (main text). All components can be either purchased from standard scientific retailers (Thorlabs, Newport), or are custom-designed, in which case we include the necessary files for producing them.

### DESIGN OF MOLECULAR CONSTRUCTS AND FUNCTIONALIZATION PROTOCOLS

#### Fluid chamber functionalization

Our fluid chamber design comprises three components: 1) a 22x22 mm top glass; 2) a laser cut parafilm spacer; 3) a 40x24 mm bottom glass (Fig. 4). Similar to the protocol described previously [1], the bottom glass is functionalized to harbor the chloro-alkane O4 HaloTag ligand (element iii, Fig. 4). This is done through a three-step protocol, that

| Number in Fig. 1 | Element |
|---|---|
| 1 | 95 mm Construction Rail, 500 mm height. XT95-500, (Thorlabs). |
| 2 | Optical Rail Carriage, M-CXL95-80, (Newport). |
| 3 | Objective, 100 X Plan Apo f=160 mm (Zeiss). |
| 4 | P-725.CDD PIFOC High Dynamics Piezo Scanner, (PI). |
| 5 | Z-flexor stage. SM1Z - Z-Axis Translation Mount (Thorlabs). |
| 6 | 50:50 Beam Splitter, BSM10R-25x36 (Thorlabs). |
| 7 | Filter Green Filter, $\lambda \sim$550 nm, (Thorlabs) |
| 8 | Cage System Iris Diaphragm, CP20S, (Thorlabs) |
| 9 | 30 mm Cage Plate with 1" Double Bore, CP35-30, (Thorlabs). |
| 10 | Plano Convex lens (Thorlabs) |
| 11 | Cage System U-Bench M6 and M4 Tapped Holes, CB1/M, (Thorlabs) (x2) |
| 12 | LED source, MCWHL5-C4-6500 K, (Thorlabs). |
| 13 | Custom cage for electronics. |
| 14 | CMOS Camera, MQ013MG-ON, (Ximea) |
| 15 | Plano-convex lens, (Thorlabs) |
| 16 | Mount-piece for tape-head (CNC fabricated). |
| 17 | Magnetic Tape Head, 902836 (Brush Industries) |
| 18 | Manipulation fork (Metal stamp, custom-made). |
| 19 | Fluid Chamber (custom-made) |
| 20 | XY Linear stage, miniature, MS-125-XY (NewPort). |
| Not shown | DAQ control card, USB-6341, (National Instruments). |
| Not shown | 12 Volts batteries, 18 Ah). |
| Not shown | Battery charger. |
| Not shown | Piezo controller (all PI) E-515.0X (chassis); E-505.00 (amplifier); E-509.01 (servo). |

TABLE I. Parts list for the magnetic tweezers setup

after cleaning the glass surface, involves its silanization to incorporate an amino-group to the glass surface (element ii, Fig. 4). After silanization, the chamber is assembled by sandwiching the three components on a hot plate with weight on them. Prior to that, the top glasses are functionalized with repel silane to make them hydrophobic (Fig. 4, a). Once the fluid chamber is assembled, the glutaldehyde group is incorporated (ii), which plays a double role: first, to allow the anchoring and immobilization of non-magnetic amine beads, used as reference beads; second, reaction with the O4 HaloTag ligand, which allows the covalent binding of HaloTagged proteins.

### Molecular Biology Protocols

All the reagents employed in this research were purchased from Sigma-Aldrich, unless otherwise specified. Both $R3^{IVV}$ and protein L monomers were cloned into and expressed from a modified pFN18a-HaloTag T7 Flexivector (Fig. 5), that contains two copies of the Spy0128 gene followed by an AviTag(pFN18a-HaloTag-TEV site-(Spy0128)2-AviTag). All protein constructs contain a 6xHisTag for purification before the AviTag. All the cloning steps were done following the strategy previously described [6]. Briefly, the genes encoding the proteins contain a 5' BamHI and a 3' BglII restriction sites. The pFN18a plasmid has a BamHI restriction site between the TEV site and the first copy of the Spy0128 gene, and the digestion with BamHI generates compatible ends with BglII. The digested gene (BamHI/BglII) encoding the protein of interest can be inserted and ligated into the dephosphorilated pFN18a (BamHI). Due to the lack of directionality, the gene can be inserted in the correct (5' to 3') or incorrect (3' to 5') orientation in the plasmid. This is checked with a subsequent digestion with BamHI and BlpI (91 bp beyond the stop codon in the 3' end). In the correct orientation, this digestion produces a fragment of 3900 bps (pFN18a-HaloTag-TEV site-BamHI) and a fragment of 1900+protein of interest sequence bp that accounts for BamHI-(protein of interest)-(Spy0128)2-AviTag-BlpI. In the case of an antisense ligation, the digestion produces the fragments pFN18a-HaloTag-TEV site-protein of interest sequence (3900 bps + sequence of gene bp) and a fragment of 1900 bp (BamHI-(Spy0128)2-AviTag-BlpI). All the cloning and amplification steps were conducted on the Escherichia coli XL10-Gold strain (Agilent Technologies). Finally, we sequence the construct to corroborate the correct insertion orientation of the gene and the fidelity of the sequence

E. coli ERL strain cells (gift of R. T. Sauer) were transformed with the plasmids containing the chimeric proteins. Protein expression inductions and purification protocols were done as previously described [7]. Briefly, the transformed cells were grown at 37 C with 250 rpm constant shaking until reached OD600 0.6. 1 mM IPTG was used to induce expression at 25 C with 250 rpm constant shaking for 12-16 h. Protein extraction from cells was done through mechanical lysis with a French press (Sim-Aminco), and the protein isolation from the lysate was done with the His60 Ni SuperflowResin (Clontech). Further purity was achieved with a size exclusion chromatography step (Superdex 200 FPLC column, GE Healthcare). Proteins were eluted in 10 mM Hepes (pH 7.2), 150 mM NaCl, 1 mM EDTA and stored with 10% v/v of glycerol at -80°C until use.

### CALCULATION OF THE FREE ENERGY LANDSCAPES

To estimate the free energy landscape, we use long time-series of the measured extension $x(t)$, and estimate a first "naive" free energy landscape as:

$$G(x) = -kT \ln p(x) - \frac{kT}{D} \int_{x_1}^{x_2} \dot{x}(x) dx, \tag{2}$$

where $p(x)$ is the probability density function of $x(t)$, $kT$ is the thermal energy, $D$ the diffusion coefficient, and $x_2 - x_1$ the upper and lower boundaries of $x(t)$. Equation (2) accounts for the equilibrium contribution as the simple Boltzmann inversion (first term), and a second non-equilibrium contribution that weights the directional derivatives of the trajectory (second term). In the case of long equilibrium dynamics, as it is our case, this second term has a negligible contribution to $G(x)$. In force spectroscopy experimental techniques, the instrument influences the molecular dynamics so that the measured trajectory $x(t)$ is a convolution of the actual molecular trajectory $\xi(t)$ and the instrumental effect $\Xi(t)$. The actual molecular free energy landscape $G(\xi)$ must be obtained by a deconvolution process.

In optical tweezers and AFM, the main instrumental effect arises from the stiffness of the force probe, either the optical trap or the AFM cantilever. In the case of magnetic tweezers, the magnetic trap stiffness is negligible, which gives rise to its intrinsic force-clamp conditions. However, the differential bead height is inferred using an image analysis algorithm that might introduce noise that is convoluted with the actual molecular dynamics. In this regard, we calculate the free energy landscape of $R3^{IVV}$ and protein L through a deconvolution process that requires to estimate the point-spread function of our image analysis algorithm. To this aim, we measure a long recording of two reference beads, not affected by the pulling force and firmly anchored to the substrate, so that their Brownian fluctuations are negligible, and estimate our point spread function from the extension histogram (Fig. 6A), which is a Gaussian distribution with a standard deviation of ~1.7 nm. Using this point spread function, we estimate the actual molecular free energy landscape by spectral deconvolution using the Jansson algorithm [8]. Figure 6B compares the "naive" free energy landscape (grey) and the deconvoluted one (black) for $R3^{IVV}$ at 8 pN. The deconvolution procedure is key to recover the actual characteristics of the free energy landscape, especially in the barrier region, which typically is more susceptible to be smeared by instrumental effects. With this procedure, we are able to fully characterize the free energy landscapes of protein L and $R3^{IVV}$ at different forces and describe the effect of the pulling force. Tables II and III show the properties of the free energy landscapes in terms of the free energy barrier from the folded $\Delta G_F^\dagger$ and unfolded $\Delta G_U^\dagger$ state, and the stiffness at the folded $\kappa_F$ and unfolded $\kappa_U$ basin, and the barrier top $\kappa_b$.

| Property | Velue |
|---|---|
| $\Delta G_F^\dagger$ | 15.3±0.2 pNnm |
| $\Delta G_U^\dagger$ | 11.1±0.4 pNnm |
| $\kappa_F$ | 0.28±0.02 pN/nm |
| $\kappa_b$ | 0.37±0.04 pN/nm |
| $\kappa_b$ | 0.08±0.01 pN/nm |

TABLE II. Properties of the free energy landscape of $R3^{IVV}$ at 8 pN.

| Property | Velue |
|---|---|
| $\Delta G_F^\dagger$ | 32.2±0.3 pNnm |
| $\Delta G_U^\dagger$ | 29.7±0.3 pNnm |
| $\kappa_F$ | 3.40±0.04 pN/nm |
| $\kappa_b$ | 7.83±0.40 pN/nm |
| $\kappa_b$ | 1.55±0.04 pN/nm |

TABLE III. Properties of the free energy landscape of protein L at 6 pN.

### EFFECT OF THE MECHANICAL PERTURBATION ON THE EVOLUTION OF A PROTEIN FREE ENERGY LANDSCAPE

To a first approximation, the effect of the pulling force on a biomolecular free energy landscape is often modeled as a linear perturbation $-F \cdot x$ that tilts the landscape towards the unfolded state. However, an unfolded protein is a polypeptide chain extended by force, and its equilibrium extension is dictated by standard polymer physics models such as the FJC. In this sense, the position of the unfolded basin displaces with force in a non-linear way. However, when the biomolecule explores only a narrow range of forces, this non-linear effects can be negligible. To test this property, we subtract here the energy landscapes for $R3^{IVV}$ and protein L and recover the effect of the pulling force on the tilt of these landscapes.

Figure 7A shows the landscape at 7.7 pN (red) and 8.3 pN (black), together with the algebraic difference (green). As shown, the difference between both landscapes is approximately linear, with a slope of -0.6 pN, in agreement with the difference in force between both landscapes. In this sense, the effect of the pulling force on the explored force range for $R3^{IVV}$ is well approximated as a linear tilt, likely due to the narrow range of forces. By contrast, in the case of protein L, the explored range of forces is much broader, and the effect of the polymer extension are readily observed in the shift of the unfolded basin position, which establishes the step-size dependence with force. Here, the subtraction of the landscapes at 4.5 pN (red) and 7.5 pN (black) has a nonlinear shape, indicating a more complex effect of the pulling force.

### CHARACTERIZATION OF THE DATA FOR THE PROTEIN INDIVIDUALS MEASURED IN FIGURE 5

We show here the properties of the molecules of R3$^{\mathrm{IVV}}$ and protein L measured for Figure 5. Figures 8 and 9 show the unfolding (blue) and refolding (red) rates as function of force. From the intersection of the fits to these data, we estimate the values of $F_{1/2}$ and $r_{1/2}$. Even not all proteins explore the full range of forces, we can estimate the coexistence forces and rates from the fits to the dependence of their rates on a few forces.

Tables IV and V show the date at which each protein was measured, the total duration of the experiment, and the values for $F_{1/2}$ and $r_{1/2}$.

| **Date** | **Total time** | $\mathbf{F_{1/2}}$ **(pN)** | $\mathbf{r_{1/2}}$ **(1/s)** |
|---|---|---|---|
| 26-Feb-2020 | 2 h 5 min | 9.1 | 1.0 |
| 25-Feb-2020 | 2 h 17 min | 8.4 | 1.4 |
| 11-Mar-2020 | 21 h 24 min | 7.9 | 0.8 |
| 25-Oct-2019 | 6 h 23 min | 10.0 | 0.7 |
| 8-Nov-2019 | 9 h 43 min | 10.2 | 1.7 |
| 14-Oct-2019 | 1 h 47 min | 8.7 | 1.0 |
| 31-Oct-2019 | 5 h 21 min | 9.0 | 1.5 |
| 21-Mar-2019 | 0 h 22 min | 8.9 | 1.4 |
| 14-Ap-2019 | 1 h 18 min | 8.4 | 1.4 |
| 16-Ap-2019 | 3 h 22 min | 7.4 | 0.7 |
| 30-Sep-2019 | 0 h 24 min | 8.4 | 0.8 |
| 30-Sep-2019 | 0 h 27 min | 9.2 | 0.7 |
| 19-Ap-2019 | 0 h 48 min | 9.1 | 0.5 |
| 19-Ap-2019 | 1 h 12 min | 8.4 | 1.6 |
| 19-Ap-2019 | 2 h 03 min | 9.5 | 2.3 |

TABLE IV. List of measured $R3^{IVV}$ molecules, indicating the date of measuring, total time of experiment, and their coexistence force $F_{1/2}$ and coexistence rate $r_{1/2}$.

| **Date** | **Total time** | $\mathbf{F_{1/2}}$ **(pN)** | $\mathbf{r_{1/2}}$ **(1/s)** |
|---|---|---|---|
| 14-Jan-2020 | 121 h 28 min | 8.2 | $5.6\times10^{-3}$ |
| 14-Jan-2020 | 118 h 12 min | 5.9 | $5.0\times10^{-2}$ |
| 21-Jan-2020 | 21 h 25 min | 7.0 | $5.0\times10^{-3}$ |
| 24-Jan-2020 | 1 h 57 min | 7.8 | $2.7\times10^{-3}$ |
| 31-Jan-2020 | 3 h 10 min | 9.2 | $7.5\times10^{-3}$ |
| 5-Feb-2020 | 41 h 45 min | 8.8 | $4.5\times10^{-3}$ |
| 5-Oct-2020 | 12 h 22 min | 8.5 | $6.0\times10^{-3}$ |
| 6-Oct-2020 | 1 h 01 min | 9.9 | $3.3\times10^{-3}$ |
| 6-Oct-2020 | 1 h 20 min | 9.5 | $3.0\times10^{-3}$ |

TABLE V. List of measured protein L molecules, indicating the date of measuring, total time of experiment, and their coexistence force $F_{1/2}$ and coexistence rate $r_{1/2}$.

*

[1] I. Popa, J. A. Rivas-Pardo, E. C. Eckels, D. J. Echelman, C. L. Badilla, J. Valle-Orero, and J. M. Fernandez, "A halotag anchored ruler for week-long studies of protein dynamics," *Journal of the American Chemical Society*, vol. 138, no. 33, pp. 10546–10553, 2016. PMID: 27409974.
[2] H. Guo, "A simple algorithm for fitting a gaussian function," *IEEE SIGNAL PROCESSING MAGAZINE*, vol. 10.1109/MSP.2011.941846, pp. 134–137, 2011.
[3] R. Tapia-Rojo, E. C. Eckels, and J. M. Fernández, "Ephemeral states in protein folding under force captured with a magnetic tweezers design," *Proceedings of the National Academy of Sciences*, vol. 116, no. 16, pp. 7873–7878, 2019.
[4] A. Alonso-Caballero, R. Tapia-Rojo, C. L. Badilla, and J. M. Fernandez, "Magnetic tweezers meets afm: ultra-stable protein dynamics across the force spectrum," *bioRxiv*, 2021.
[5] R. Tapia-Rojo, Á. Alonso-Caballero, and J. M. Fernández, "Talin folding as the tuning fork of cellular mechanotransduction," *Proceedings of the National Academy of Sciences*, vol. 117, no. 35, pp. 21346–21353, 2020.
[6] M. Carrion-Vazquez, A. F. Oberhauser, S. B. Fowler, P. E. Marszalek, S. E. Broedel, J. Clarke, and J. M. Fernandez, "Mechanical and chemical unfolding of a single protein: A comparison," *Proceedings of the National Academy of Sciences*, vol. 96, no. 7, pp. 3694–3699, 1999.
[7] J. Alegre-Cebollada, C. L. Badilla, and J. M. Fernández, "Isopeptide bonds block the mechanical extension of pili in pathogenic streptococcus pyogenes," *Journal of Biological Chemistry*, vol. 285, no. 15, pp. 11235–11242, 2010.
[8] P. A. Jansson, "Deconvolution of images and spectra," 1997.

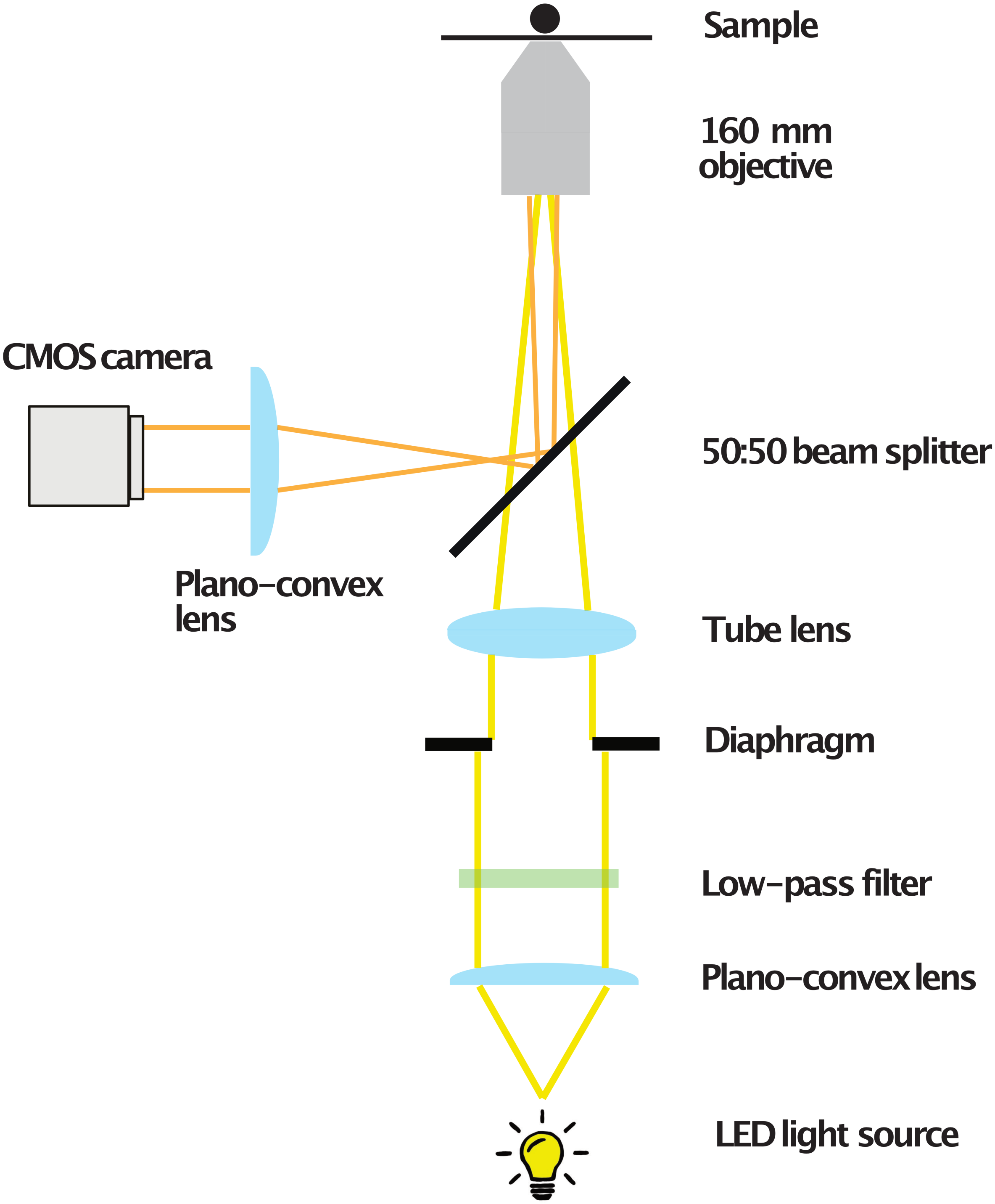


FIG. 1. **Schematics of the optical path**: Light (yellow beam) from a LED source is focused to a 160 mm objective using two plano convex lenses. The image (orange beam) is directed to a camera using a 50:50 beam splitter and plano convex lens to convert the 160 mm beam to a parallel one.

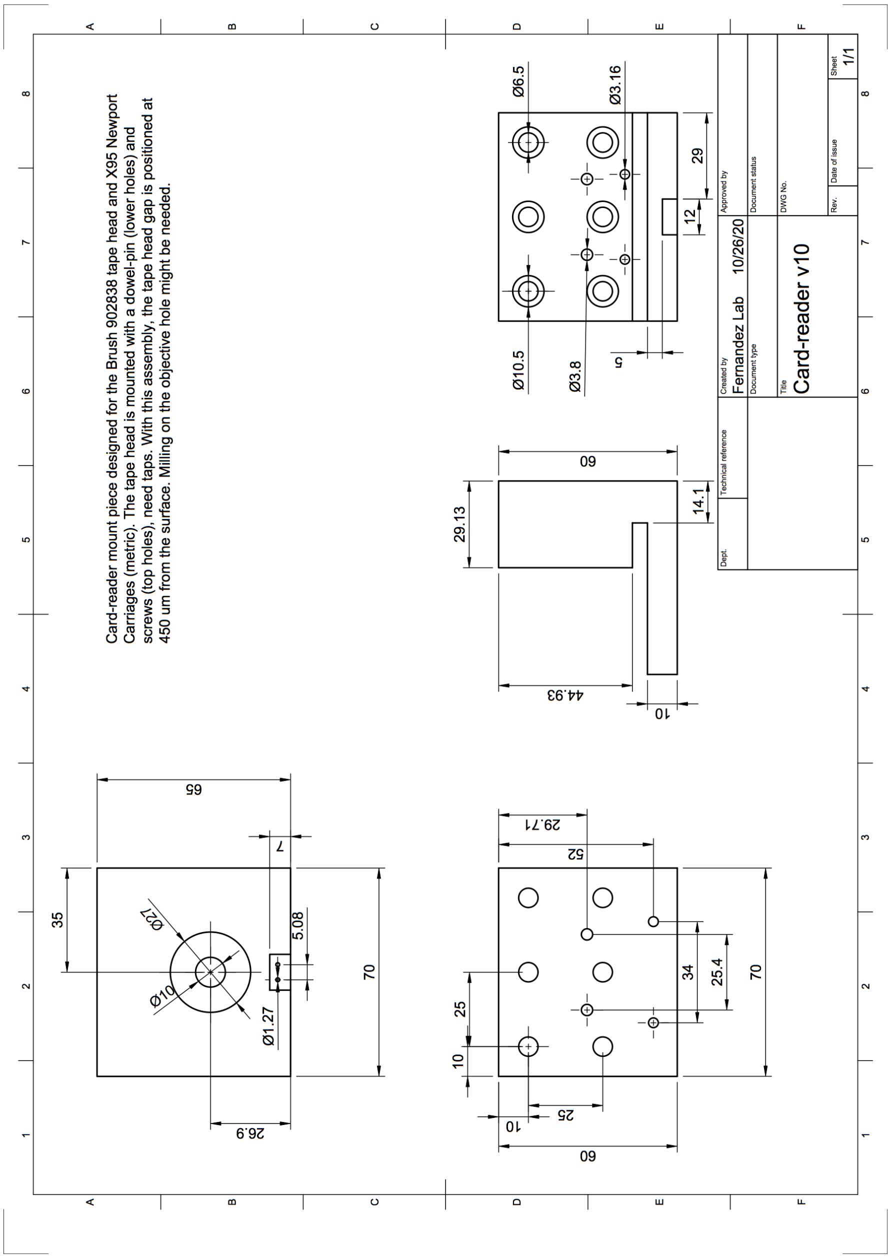


FIG. 2. Technical drawing of the tape head mounting piece

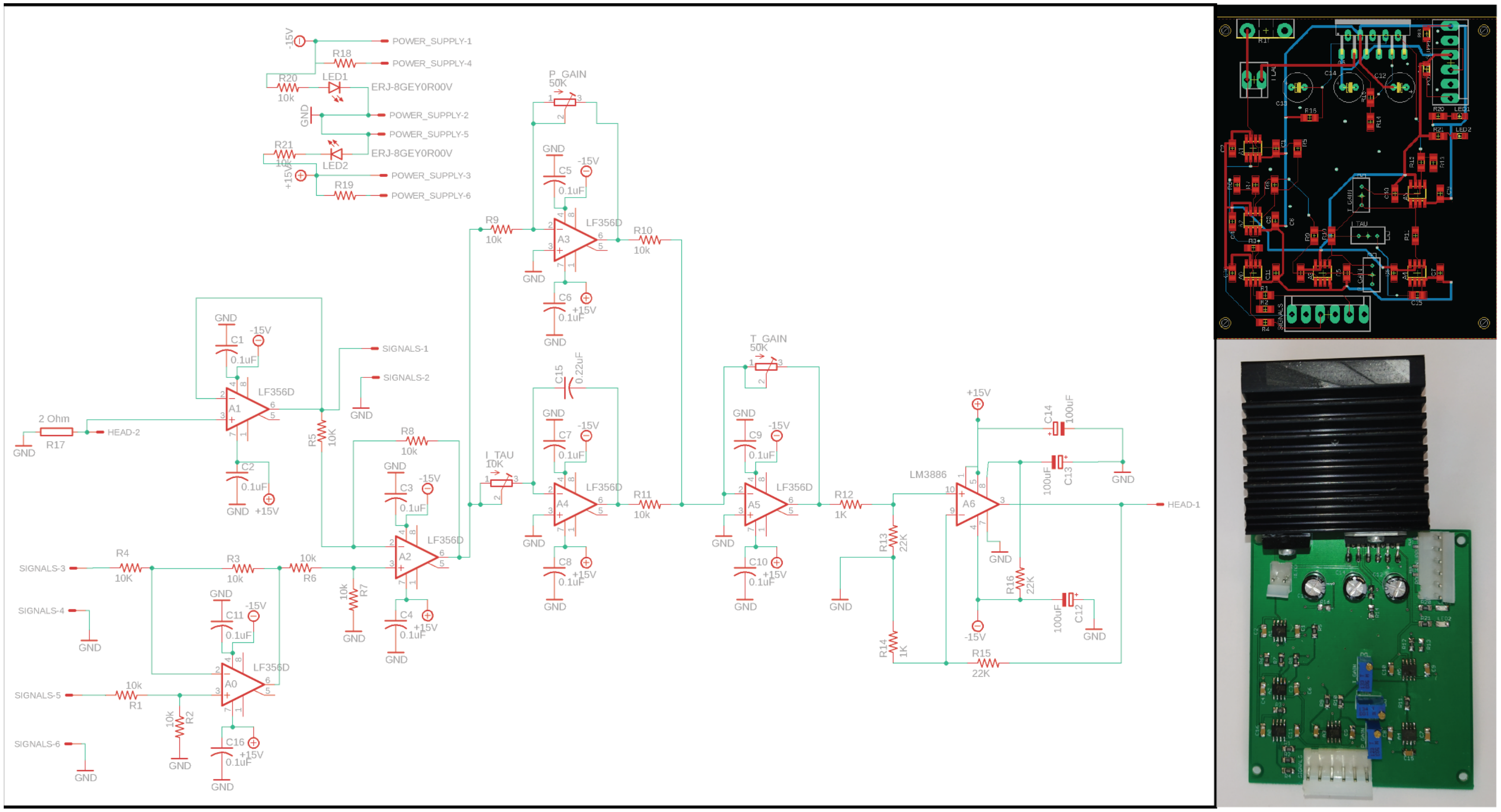

FIG. 3. **PI Control Circuit of the Magnetic Tape Head.** (Left) Schematic diagram of the PI circuit design. (right, up) Layout for the board for surface mount. (right, down) Picture of the assembled circuit. Eagle file included as SI File.

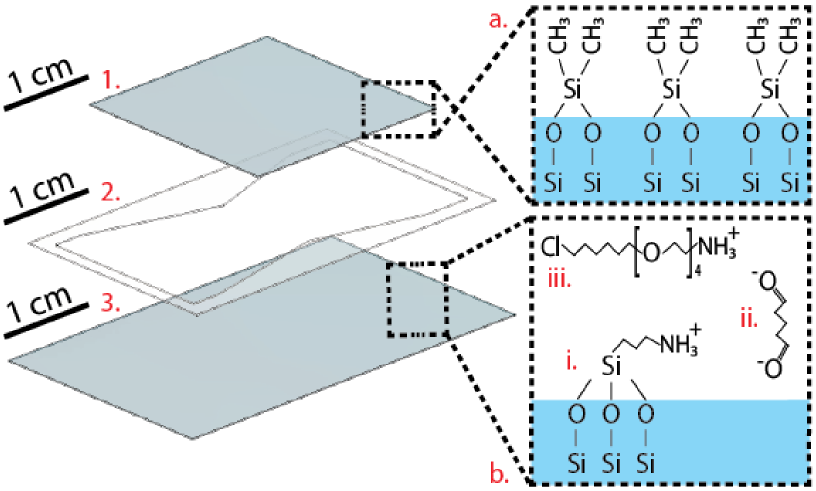


FIG. 4. **Schematics of the fluid chamber assembly and manipulation**

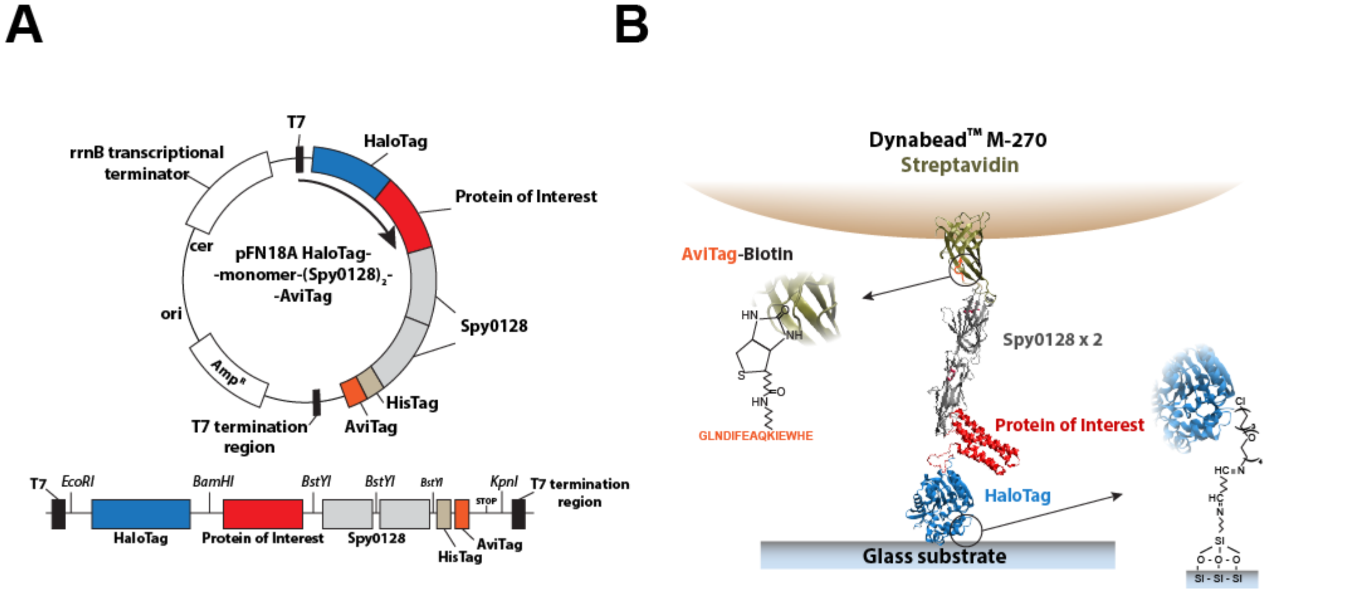


FIG. 5. **Expression vectors and single-molecule magnetic tweezers tethering.** A) Modified pFN18a HaloTag T7 Flexi vector. The T7 promoter regulates the expression of the protein HaloTag-monomer(here, talin R3 domain IVVI)-(Spy0128)2-HisTag-AviTag. Below is shown the 5'-3' scheme of the expression cassette with the restriction enzyme sites formed after subcloning. B) Magnetic tweezers single-molecule tethering. The N-terminal HaloTag binds covalently to a glass surface functionalized with the HaloTag ligand. The HaloTag ligand is covalently crosslinked to the surface through a glutaradehyde molecule that bridges the amino group of the ligand with the amino group of an APTES molecule. On the C-terminus of the protein, the biotinylated AviTag is recognized by the streptavidin molecules that coat the superparamagnetic bead (DynabeadM-270 Streptavidin). The two Spy0128 modules act as an inextensible molecular spacer that minimizes unspecific bead-surface interactions. Here the protein of interest is the R3$^{IVV}$ domain from mouse talin-1, but any single domain protein or polyprotein can be inserted in this vector.

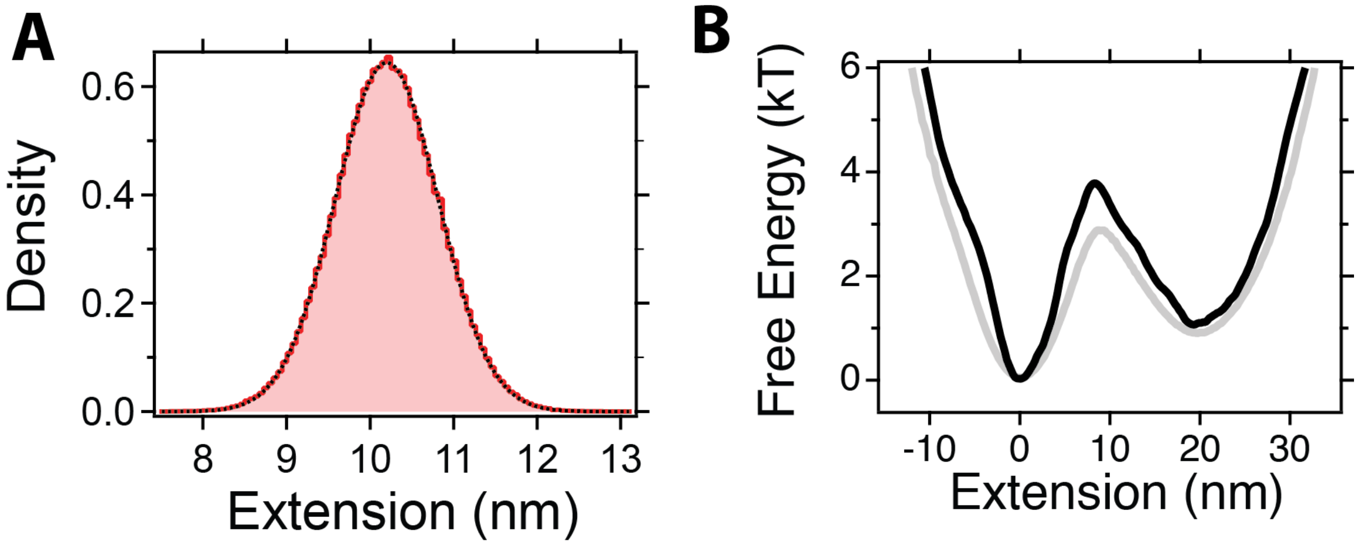


FIG. 6. **Point-spread-function:**

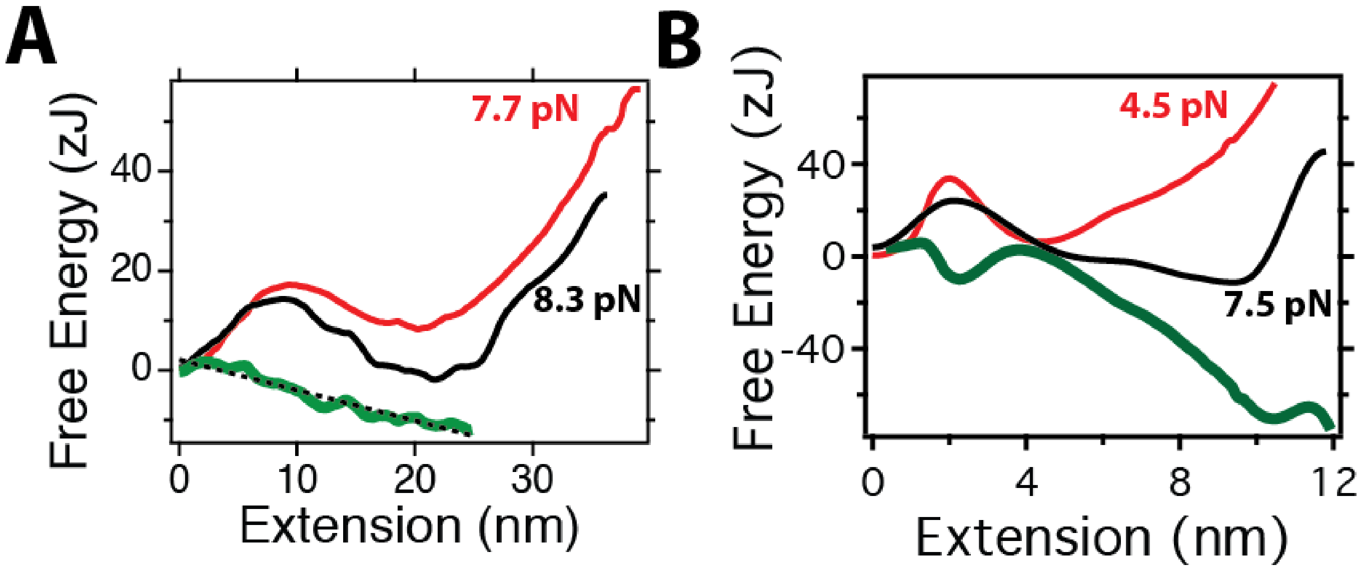


FIG. 7. **Difference between the free energy landscapes at different forces to infer the mechanical perturbation**. (A) Difference in the landscapes at 7.7 pN (red) and 8.3 pN (black) for R3$^{IVV}$. The mechanical perturbation has roughly linear dependence, which a slope of ∼0.6 pN, which successfully recovers the perturbation between both landscapes. (B) Difference in the landscapes at 4.5 pN (red) and 7.5 pN (black) for protein L. The difference between both landscapes has a complex non-linear form, that indicates that the mechanical perturbation cannot be understood here with a simple $-Fx$ effect.

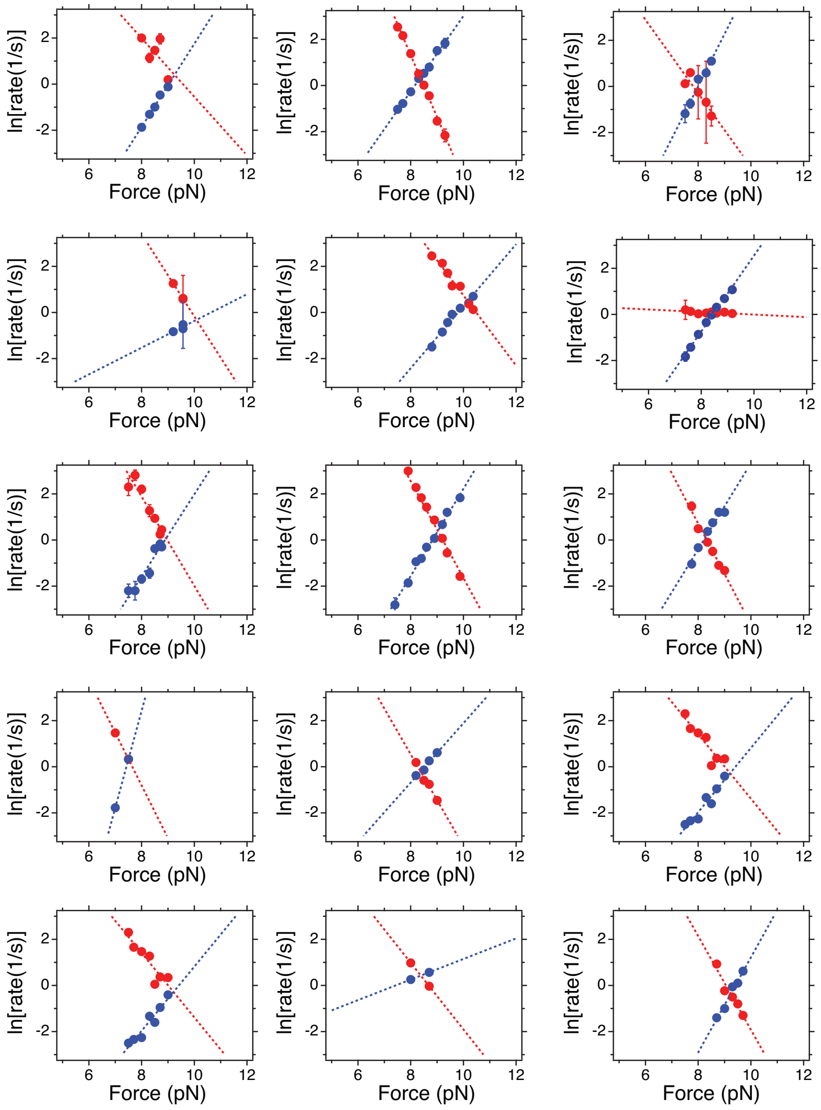


FIG. 8. **Folding (red) and unfolding rates for the measured R3$^{IVV}$ molecules**: The values of $F_{1/2}$ and $r_{1/2}$ employed for Fig. 4 (main text) are obtained from the intersection of the fits to this data (dotted lines).

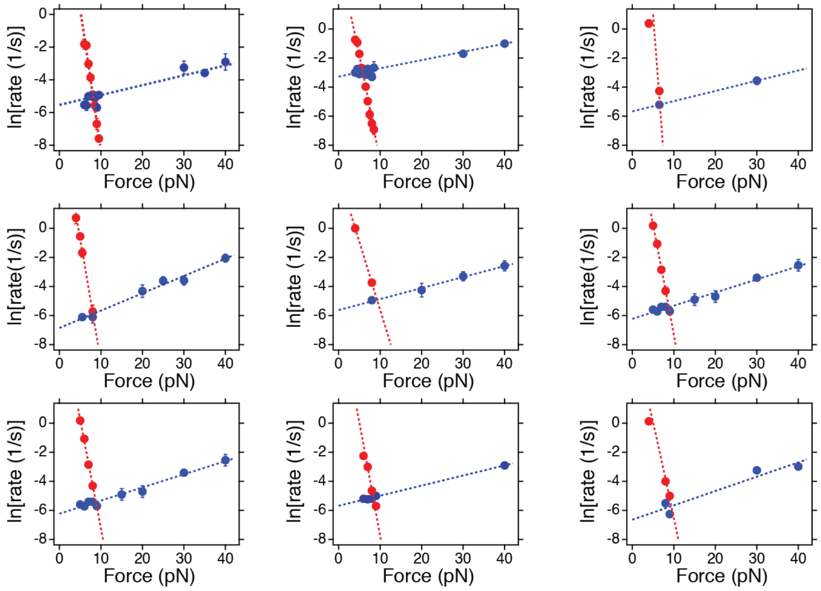


FIG. 9. **Folding (red) and unfolding rates for the measured protein L molecules**: The values of $F_{1/2}$ and $r_{1/2}$ employed for Fig. 4 (main text) are obtained from the intersection of the fits to this data (dotted lines).
